# Modeling Abiotic Niches of Crops and Wild Ancestors Using Deep Learning: A Generalized Approach

**DOI:** 10.1101/826347

**Authors:** W. G. Hulleman, R. A. Vos

## Abstract

**Introduction:** Understanding what interactions and environmental factors shape the geographic distribution of species is one of the fundamental questions in ecology and evolution. Insofar as the focus is on agriculturally important species, insight into this is also of applied importance. Species Distribution Modeling (SDM) comprises a spectrum of approaches for establishing correlative models of species (co-)occurrences and geospatial patterns of abiotic environmental variables.

**Methods:** Here, we contribute to this field by presenting a generalized approach for SDM that utilizes deep learning, which offers some improvements over current methods, and by presenting a case study on the habitat suitability of staple crops and their wild ancestors. The approach we present is implemented in a reusable software toolkit, which we apply to an extensive data set of geo-referenced occurrence records for 52 species and 59 GIS layers. We compare the habitat suitability projections for selected, major crop species with the actual extent of their current cultivation.

**Results:** Our results show that the approach yields especially plausible projections for species with large numbers of occurrences (>500). For the analysis of such data sets, the toolkit provides a convenient interface for using deep neural networks in SDM, a relatively novel application of deep learning. The toolkit, the data, and the results are available as open source / open access packages.

**Conclusions:** Species Distribution Modeling with deep learning is a promising avenue for method development. The niche projections that can be produced are plausible, and the general approach provides great flexibility for incorporating additional data such as species interactions.

## Introduction

Understanding what processes and environmental factors shape the distribution of species still remains poorly understood. What constitutes a suitable habitat is only partially known or even completely unknown for many uncommon species. Furthermore, a number of factors including human activities like land transformation and climate change have put Earth’s biodiversity under stress [1], causing geographical distributions to shift [2][3] or change through degradation of habitat suitability.

Aside from the current biodiversity loss, these changes in land use and climate are also highly relevant to agriculture, as climate is an important determinant for agricultural productivity [4]. As such, changes in climate or human activities that alter the environment have the potential to cause a decrease in agricultural yield and lead to economic losses [5]. To gain insight into the current habitat suitability of a variety of crop species, this study will model the global habitat suitability for a number of crops and their ancestors. These results paired with predictions on future habitat suitability could mitigate economic losses and aid global food security in the future.

However, a number of challenges can arise when predicting the distribution of species. A paper by Hickisch et al. [6] elaborates several reasons that affect the availability of (spatial) biodiversity data. First, there are geographic and taxonomic sampling biases that can make it harder to obtain data on certain groups of species (e.g. species that are hard to find or identify). Some species are under-sampled as locations are physically harder to reach. Other reasons that can contribute to inconsistent spatial data include policy bias, where more samples are recorded in areas where conservation efforts are focused. Considering these biases, using limited and/or geographically biased data can have implications for the reliability of any predictions about the presence or absence of a species [7].

Since 1980 several approaches have been used to model the realized, abiotic niches of species (commonly called ecological niche models, ENMS; or species distribution models, SDMs). The number of available methods, and their accuracy has gradually increased over time and has led to a widespread use of SDMs by the 2000s [8]. An SDM can be defined as follows: ‘a numerical tool that combines observations of species occurrence or abundance with environmental estimates’ and they utilize this information to predict potential distribution, or habitat suitability, by extrapolation [8].

Approaches to creating SDMs are numerous; the chosen approach needs to be carefully considered depending on the type of data that is available and the species it is applied to. Approaches include methods such as generalized linear models (GLMs), generalized adaptive models (GAMs), random forest, support vector machines, maximum entropy, and artificial neural networks (ANNs) [9][10]. MaxEnt [11] is a broadly used software environment for creating SDMs, based on a maximum entropy approach. Its popularity is evident from the over 1000 published applications between its introduction in 2006 and a later review by Merow et al. in 2013 [12]. This is due to its superior predictive accuracy when compared to other approaches and its ease of use, allowing users without any programming experience to create complex SDMs [12].

However, since the introduction of MaxEnt in 2006, artificial neural networks (ANNs) - and specifically deep learning - have been shown (e.g. by Rademaker et al. [13] and Botella et al. [14]) to achieve similar accuracies to the maximum entropy approach. Deep learning offers the advantage of using the presence of species as additional variables, allowing co-occurrence of species with which the focal species interacts, or those with similar ecological niches, to be used to predict habitat suitability for a species of interest. This entails that the model is able to make use of biotic variables (i.e. variables that represent living organisms), in addition to the more traditional abiotic variables (variables that represent intimate features or processes in the ecosystem). A direct comparison between a deep learning and maximum entropy approach reveals that a deep learning SDM is more accurate at predicting potential distribution if the species in question has a substantial number of occurrences to be trained on. Nevertheless, deep learning SDMs have shown to perform worse with low numbers of occurrences [13].

Another drawback of using deep learning-based models for the projection of potential species distribution is lack of user-friendliness. Whereas MaxEnt offers open source software that uses a GUI to guide people through the process of model creation, any deep learning model will require the user to build such a model from scratch, in code. This means that any user without experience in programming, modeling and deep learning is unable to fully use its potential. Here, this challenge is addressed by presenting a framework for the construction and evaluation of SDMs with deep learning (hereafter: ‘sdmdl’) implemented in the python programming language. Some of the tangible results presented here thus constitute a software package called ‘sdmdl’ that automates the creation of species distribution models. This package was developed to meet the following requirements:

- Maximizing ease of use by only requiring from the user a few basic commands.
- Maximizing the package’s flexibility to optionally change parameters of the model.
- Testable functionality of the package by way of unit tests.
- Demonstrable utility by applying the resulting package to a case study.
- A clear outlook on future improvements and feature implementations.

To demonstrate the utility of this package we apply it to model and project habitat suitability worldwide for a number of staple crops and their wild relatives.

## Methods

### Software development

All software development and testing for the package ‘sdmdl’ was performed using python version 3.6 [15]. The main Integrated Development Environment (IDE) used to write the source code of the package was PyCharm Free Community [16], although Spyder [17] was used during the initial phases of development. The source code, required input files, documentation and all testing code that were created during development are available through a GitHub repository (https://github.com/naturalis/sdmdl). Details on how to install and use the package are provided at: https://sdmdl.readthedocs.io.

One of the goals of this study was that the code of the created package was testable so as to guarantee reproducibility of its functionality. This has been performed using python’s unit testing framework. To ensure that code would be repeatable at all times, continuous integrating provided by Travis CI (see https://travis-ci.com/naturalis/sdmdl) was used. This is a service that tests the code of a GitHub repository every time code is modified or new code is added. Using a set of predefined instructions, Travis CI creates a remote virtual Linux environment to perform all of the following procedures automatically:

1. Install a specific version of a requested programming language, in the case of this study this was Python 3.6.
2. Install any non-python software dependencies (e.g. GDAL for spatial computation).
3. Install all python package dependencies for the created package.
4. Install and run pytest, which automatically tests package code by running all scripts which file name start with ‘test_’.
5. Once the tests have been successfully completed returns the testing coverage.

### Case study preliminaries

The created package and its functionality are assessed and demonstrated by way of the case study, outlined in the Introduction, of crops and their wild ancestors. The occurrence data collected for this case study are available in unprocessed format as doi:10.5281/zenodo.3460512 and in preprocessed format as doi:10.5281/zenodo.3460530. A list of the environmental variables that were used to create the case study models can be found in the Supplementary Table, and can be obtained from doi: 10.5281/zenodo.3460541. The following two subsections describe the preliminary work to collect and clean the data used for the case study.

#### Literature review

Due to the nature of the case study, literature review was required before the occurrence data could be obtained for each species. This initial literature review was performed for a list of species based on major crop types in the standard cross cultural sample [18]. This resulted in a list of species that are considered the closest genetic wild ancestors (see Table 1). In most cases there is one ancestor per crop, although sometimes multiple ancestors can be identified in the case of complex hybridization events. Accompanied by a bibliography with the salient literature on the domestication of each of the crops.

**Table 1:** Results of literature research. For each crop, the list displays the scientific name of its closest ancestor(s), references to literature supporting their genetic relation and geographical distribution. Also shows the number of occurrences per species after the data has been cleaned (see the occurrence coordinate cleaning section in methods). References in this table are numbered separately from the other references in this paper, the references for this table can be found in the Supplementary References.

| **Crop species** | **Scientific Name** | **References\*** | **Occurrences** |
| --- | --- | --- | --- |
| Barley | *Hordeum vulgare* |  | 14919 |
|  | *Hordeum spontaneum* | [1][2] | 2720 |
| Breadfruit | *Artocarpus altilis* |  | 457 |
|  | *Artocarpus camansi* | [3][4] | 9 |
| Broom-corn | *Sorghum bicolor* |  | 11318 |
|  | *Sorghum bicolor* subsp. *verticilliflorum* | [5][6] | 154 |
| Cassava | *Manihot esculenta* |  | 3607 |
|  | *Manihot esculenta* subsp. *flabellifolia* | [7][8] | 364 |
| Common wheat | *Triticum aestivum* |  | 11572 |
|  | *Triticum urartu* | [9][10] | 278 |
|  | *Aegilops speltoides* | [9][10] | 398 |
|  | *Triticum turgidum* subsp. *dicoccoides* | [9][10] | 743 |
|  | *Aegilops tauschii* | [9][10] | 800 |
| Cushaw pumpkin | *Cucurbita argyrosperma* |  | 803 |
|  | *Cucurbita argyrosperma* subsp. *sororia* | [11][12] | 350 |
| Dates | *Phoenix dactylifera* |  | 1145 |
|  | *Phoenix sylvestris* | [13][14][15] | 6 |
| Finger millet | *Eleusine coracana* |  | 1634 |
|  | *Eleusine africana* | [16][17] | 357 |
| Maize | *Zea mays* |  | 15495 |
|  | *Zea mays* subsp. *mexicana* | [18][19][20] | 604 |
|  | *Zea mays* subsp. *parviglumis* | [18][19][20] | 617 |
| Mango | *Mangifera indica* |  | 2996 |
|  | *Mangifera sylvatica* | [21][22] | 9 |
| Peanut | *Arachis hypogaea* |  | 4173 |
|  | *Arachis duranensis* | [23][24] | 283 |

Table 1: Continue
| Crop species | Scientific Name | References* | Occurrences |
| --- | --- | --- | --- |
| Plantain | *Musa x paradisiaca* |  | 429 |
|  | *Musa acuminata* | [25][26] | 181 |
|  | *Musa balbisiana* | [25][26] | 100 |
| Potato | *Solanum tuberosum* |  | 7862 |
|  | *Solanum bukasovii* | [27][28] | 181 |
| Purple yam | *Dioscorea alata* |  | 713 |
|  | *Dioscorea hamiltonii* | [29] | 95 |
|  | *Dioscorea persimilis* | [29] | 75 |
| Rice | *Oryza sativa* |  | 7145 |
|  | *Oryza rufipogon* | [30][31] | 3144 |
| Rye | *Secale cereale* |  | 7744 |
|  | *Secale vavilovii* | [32][33] | 10 |
| Squash | *Cucurbita maxima* |  | 1249 |
|  | *Cucurbita maxima* subsp. *andreana* | [11] | 28 |
| Sweet potato | *Ipomoea batatas* |  | 3877 |
|  | *Ipomoea trifida* | [34][35] | 904 |
| Taro | *Colocasia esculenta* |  | 1512 |
|  | *Colocasia formosana* | [36][37] | 23 |
| Teff | *Eragrostis tef* |  | 287 |
|  | *Eragrostis pilosa* | [38][39][40] | 3639 |
| White yam | *Dioscorea rotundata* |  | 346 |
|  | *Dioscorea cayennensis* | [41][42] | 474 |
|  | *Dioscorea praehensilis* | [41][42] | 307 |
|  | *Dioscorea abyssinica* | [41][42] | 135 |
| Yellow yam | *Dioscorea cayennensis* |  | 582 |
|  | *Dioscorea minutiflora* | [43] | 167 |

Once this review was completed, a small secondary review was performed to identify the native range of each wild ancestor. These distributions were used to spatially query the occurrence data of the wild relatives, so only occurrences from its native range would be included.

#### Occurrence data collection

Once these reviews were completed the occurrence data was obtained for each of the crops and their closest wild relative(s). Occurrence archives downloaded from the Global Biodiversity Information Facility (or GBIF) database [19] contain an occurrence table and a number of metadata files in DarwinCore archive format [20]. For the purpose of this study the only required data was the occurrence table for each species, from which the following four columns were extracted for each occurrence: (1) gbifID, i.e. a unique identifier for each occurrence in the GBIF database (2) decimalLatitude, the (decimal) latitude value of an occurrence (3) decimalLongitude, the (decimal) longitude value of an occurrence (4) acceptedScientificName, the scientific species name of the occurrence as considered ‘accepted’ by the GBIF taxonomic backbone.

#### Occurrence coordinate cleaning

To use the occurrence coordinates for training a model they first needed to be cleaned. This involved removing any invalid data that could potentially cause problems during the data preparations or model training, and removing any occurrences that have spatial issues. This was done using the ‘CoordinateCleaner’ [21] package in R [22]. To preprocess the occurrence datasets used in the case study a pipeline involving eleven steps was followed, sequentially removing occurrences if:

1. The precision of the latitude or longitude coordinate is less than 2 decimal values.
2. The latitude and longitude fall outside the valid range, which here refers to the size of the raster layers used as input to the model. (maximum latitude = 90, minimum latitude = - 60, maximum longitude = 180, minimum longitude = −180)
3. The exact location is not unique.
4. The occurrence is in, or nearby, country capitals.
5. The occurrence is in, or nearby, country or province centroids.
6. The occurrence has identical latitude and longitude values.
7. The occurrence is in, or near, GBIF headquarters.
8. The occurrence is in, or near, biodiversity institutions.
9. The occurrence has non-terrestrial coordinates.
10. The latitude and/or longitude is exactly 0.
11. An occurrence is geographically isolated and the distance to any other occurrence is at least 1000 kilometers.

Once these filtering steps were executed for each species, the occurrence data was ready to be consumed by the package.

### ‘sdmdl’ analysis

Here, the practical procedures that were followed to install and configure the package are omitted. A guide on how to setup and work with the created package is available at https://sdmdl.readthedocs.io. Briefly, to use the ‘sdmdl’ package, the following prerequisites needed to be met:

1. At least one occurrence table for a species of choice was provided. Here, 117,020 occurrences for 52 species were provided.
2. At least one environmental raster layer for a variable of choice was provided. Here, 59 environmental raster layers were used.
3. A copy or clone of the GitHub repository is present on the user’s device.

Subsequently, two data-generating steps were taken. Firstly, presence (i.e. co-occurrence) maps were created to be used by the model as additional variables. Secondly, pseudo-absence locations were generated by random sampling. These were used to train the model to detect which environmental variables make a location suitable or unsuitable for a given species.

#### Creating presence maps

Presence maps are raster maps where pixel value 0 represents the absence of a species and 1 represents its presence, or occurrence. These maps were created for every species in the case study, to serve as input variables to the model to help predict the presence of another, focal, species. Thus, all presence layers were used – as potential data on co-occurrence - during the modeling for a species except for the presence map of the species itself. The presence of other species served as variables that may be interpreted as an interaction or ecological niche similarity between different species. This may be applicable as the presence of a species is not always strictly based on their abiotic surroundings, it can also be dictated by the presence of other species within an ecosystem [23]. The presence maps were created by following these steps:

1. Create a copy of an empty land map (included in the GitHub repository, see Figure 1). This raster map has two distinct values: no data for any non-terrestrial locations and zero for terrestrial locations.
2. Iterate over every occurrence of a species and set the value of the occurrence locations to 1, which represents presence.
3. Repeat for all species in the set. The end result is a map marking the location of every occurrence. See Figure 2 and Figure 3 for two examples of presence maps.

**Figure 1:**
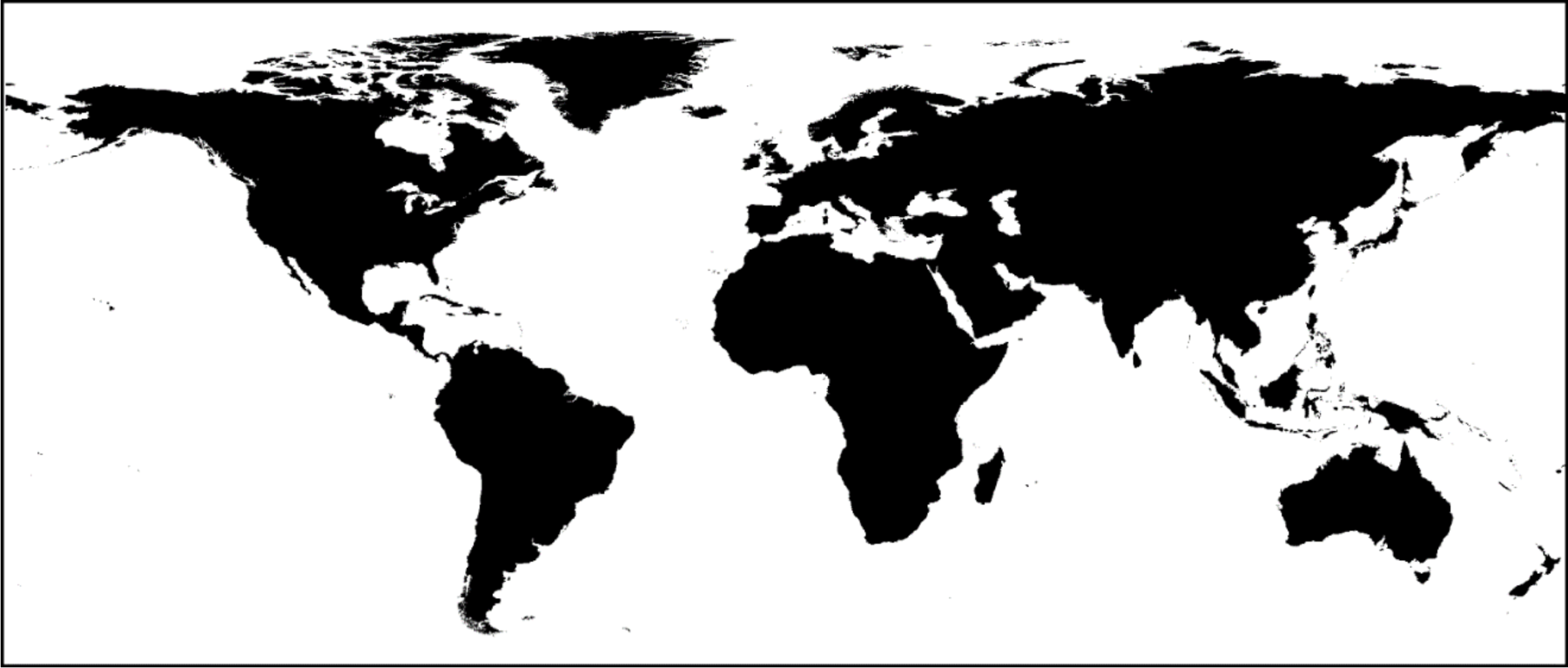
Visual representation of the empty world map.

**Figure 2:**
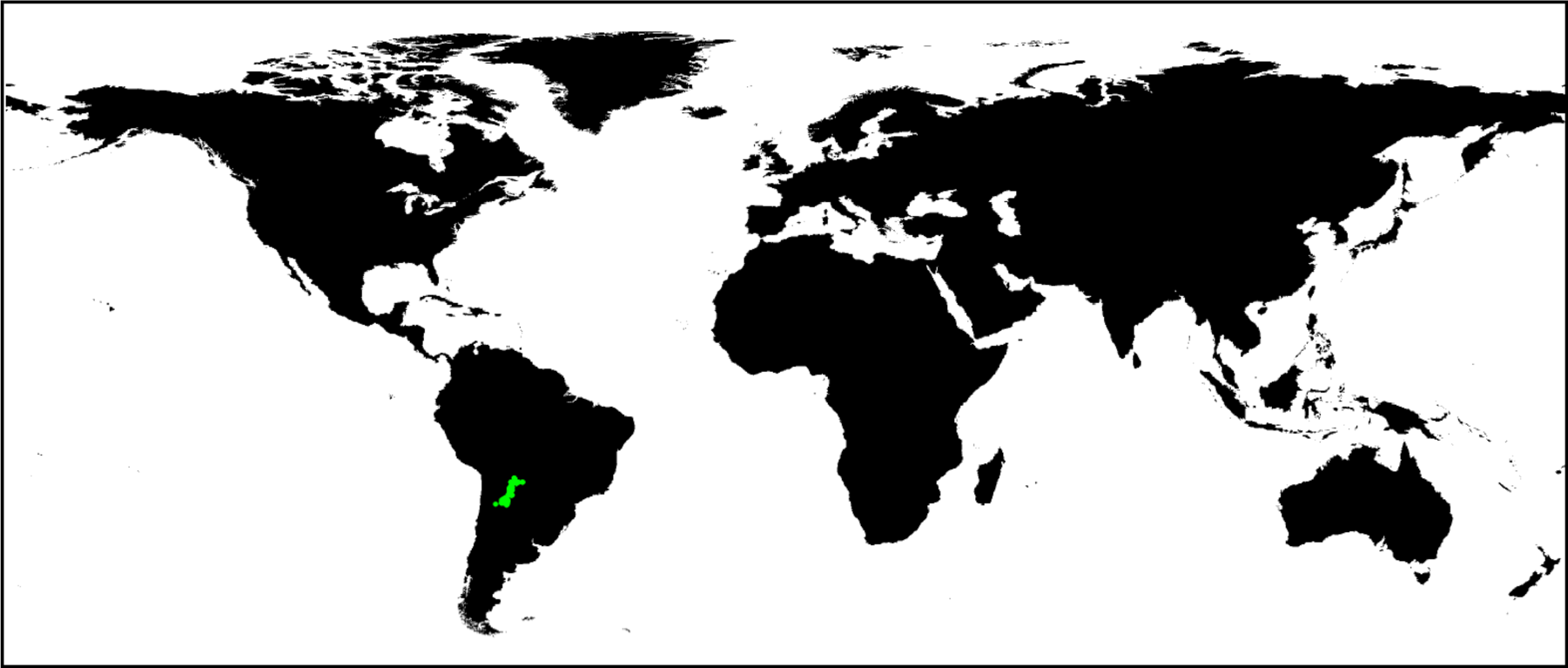
Occurrences for Arachis durnensis (n=283)

**Figure 3:**
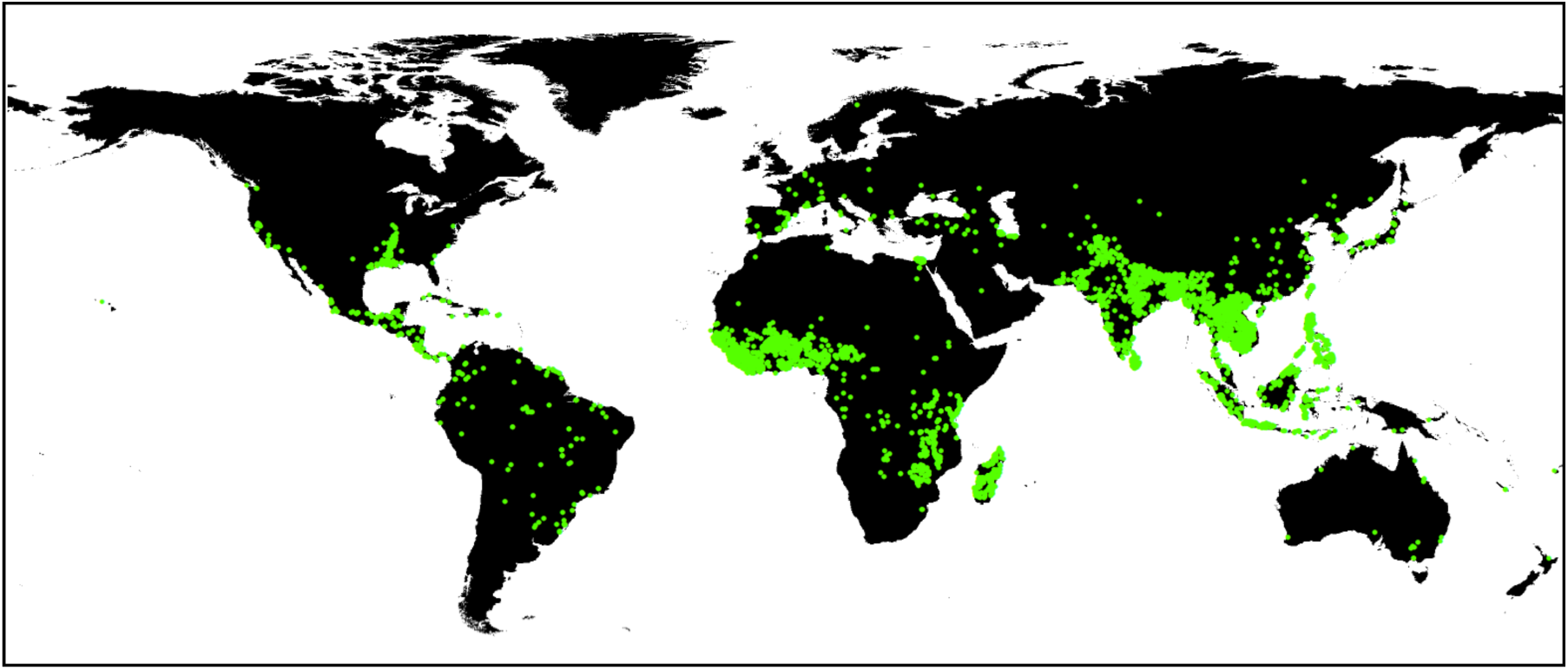
Occurrences for Oryza sativa (n=7,145)

#### Generating pseudo-absences

To correctly train a model to predict the presence of a species, it needs to be aware of what makes a location suitable or unsuitable for it. A pragmatic way to achieve this is by randomly sampling absence locations and adding these ‘pseudo-absences’ to the occurrences. By not only training the model on the locations the species occurs in, but also considering what locations are unsuitable for it, the final model can predict a species’ distribution more accurately.

The pseudo-absences were sampled according to the amount of occurrence data for the focal species. If the species had fewer than 2,000 occurrences, 2,000 pseudo-absences were sampled randomly anywhere on (terrestrial) earth (see Figure 4). If the species had more than 2,000 occurrences, the number of pseudo-absences that was sampled was equal to the number of occurrences for that species (see Figure 5). For the case study, the value 2,000 was used as the minimum number of pseudo-absences, however in other cases this value can be defined by the user using a configuration file.

**Figure 4:**
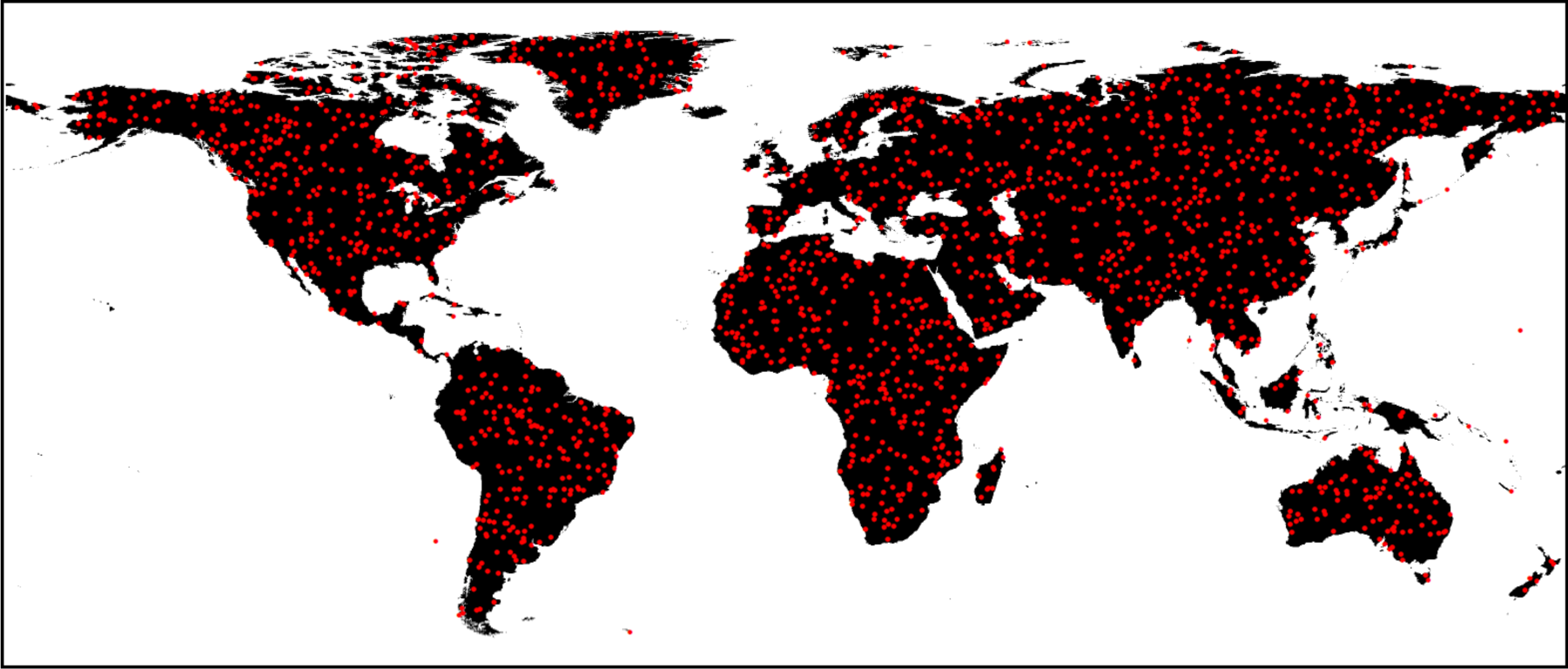
Pseudo-absences sampled for Arachis duranensis (n_occurrences_=283, n_pseudo-absences_=2,000)

**Figure 5:**
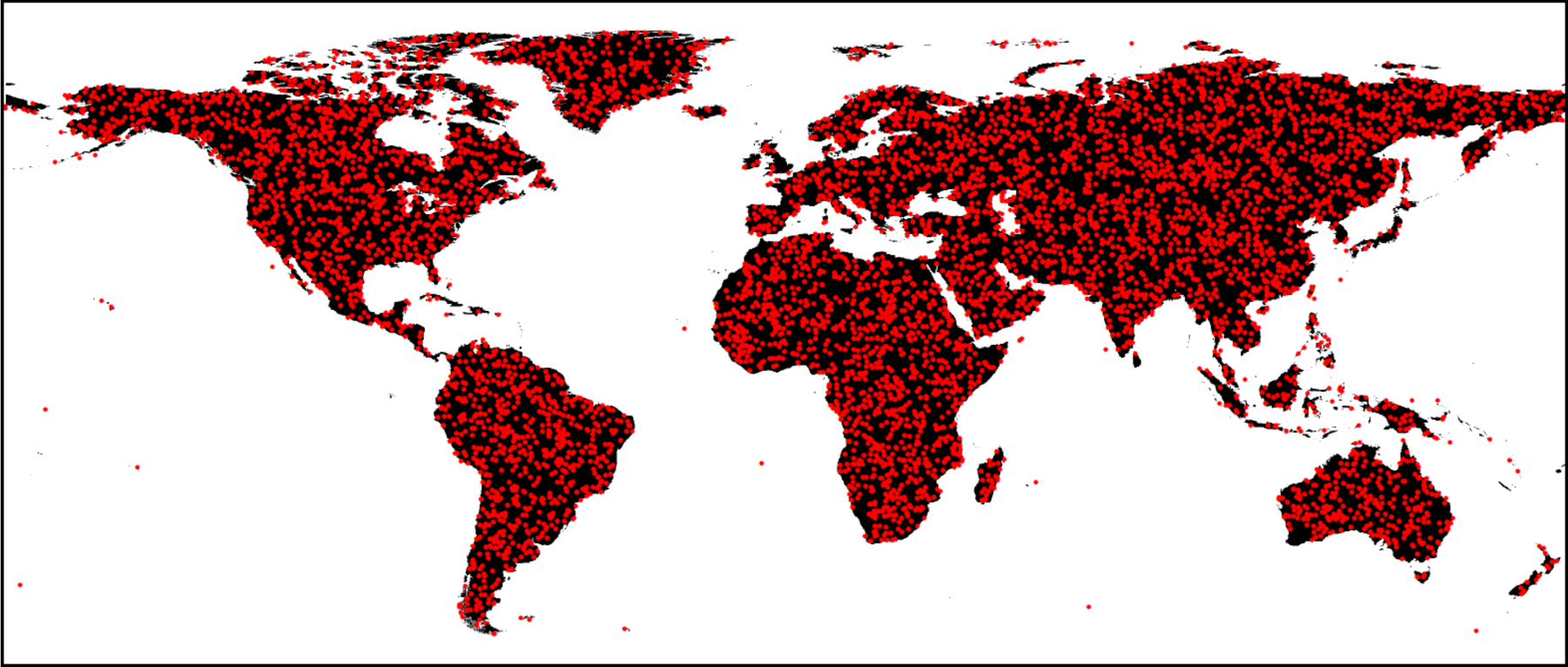
Pseudo-absences sampled for Oryza sativa (n_occurrences_=7,145, n_pseudo-absences_=7,145)

#### Preparing training data vectors

Once the two previous steps were completed, the training data vectors to the model were created. Preparing these was a process that consumed two types of data. First, the environmental raster layers collected for the case study, which were combined with the generated raster layers of the presence maps. Second, the occurrence data that were collected and cleaned, which were combined with a set of randomly sampled pseudo-absences.

The input data to the model was prepared by querying the locations of all occurrences and pseudo-absences, and obtaining the raster values for that specific location (an example of this procedure is shown in Figure 6). If a raster layer contained categorical data, the values were extracted directly; if the raster layer contained numerical data the values were scaled first using the following formula:

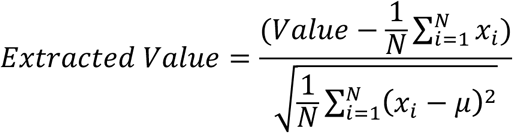

**Figure 6:**
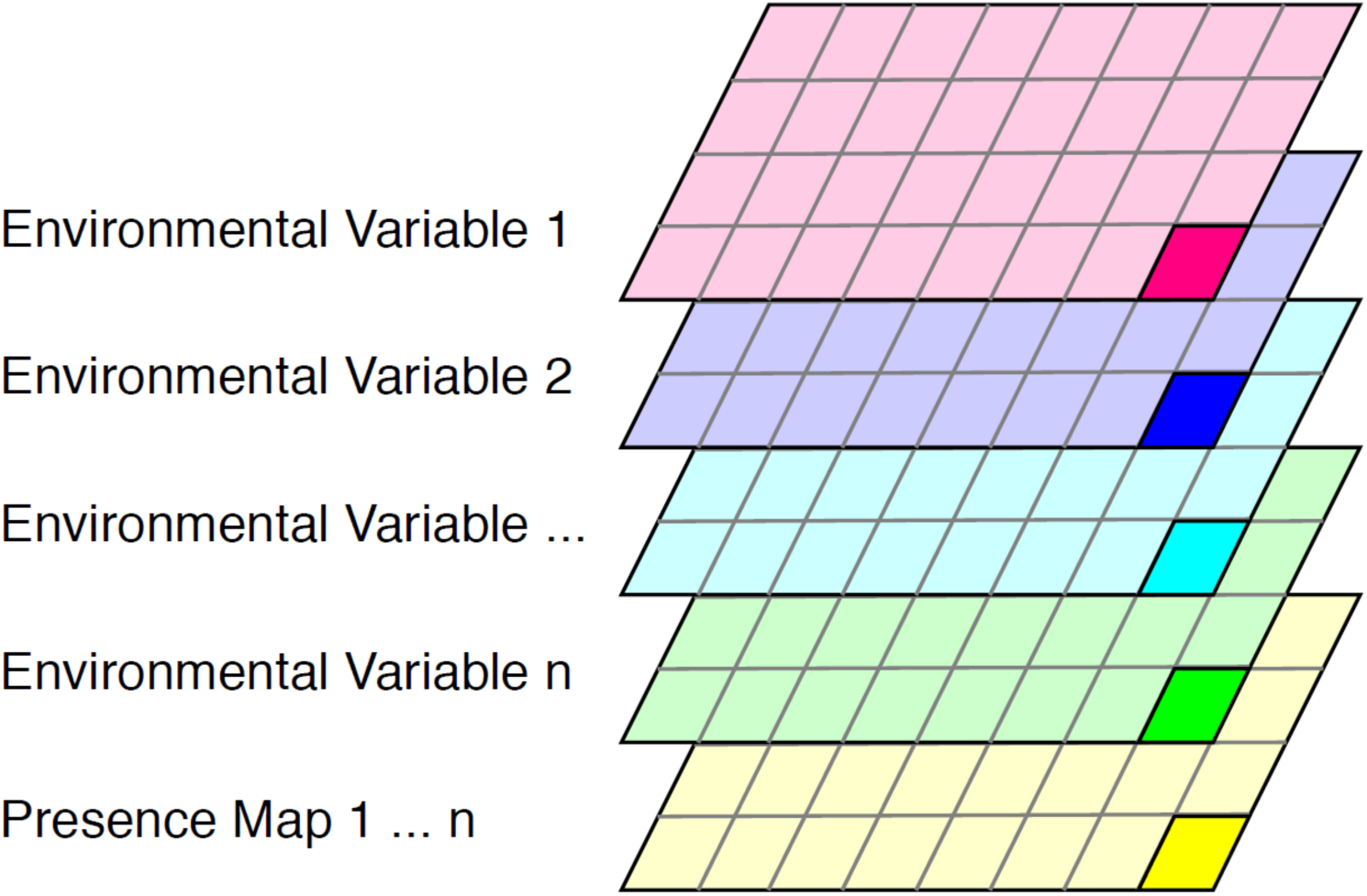
Representation of data preparation process. Each colored layer represents an environmental raster layer provided by the user, or a presence map created by the package. The highlighted pixel represents a location where an occurrence or pseudo-absence exists. At each of these locations the raster values are extracted.

#### DNN analysis

The preceding data preparation steps are broadly applicable to any method for species distribution modeling that uses some form of machine learning (including maximum entropy). The following sections describe how these data were used in an analysis that uses deep neural networks (DNNs). These sections thus include defining a DNN architecture, training it, evaluating the trained model(s) and assessing feature importance, and finally using the optimal model to predict species’ potential distributions.

#### Creating a DNN architecture

When creating and training a deep learning model there are multiple choices that can affect the performance of the final model. The first and arguably one of the most important parts of this process is creating a model architecture. Deep learning models can come in a range of different shapes and sizes. In principle, all deep learning networks have the same basic structure, which is illustrated in Figure 7. A deep learning model has a number of inputs (visualized in green), at least two hidden layers (visualized in blue) and outputs (visualized in red).

**Figure 7:**
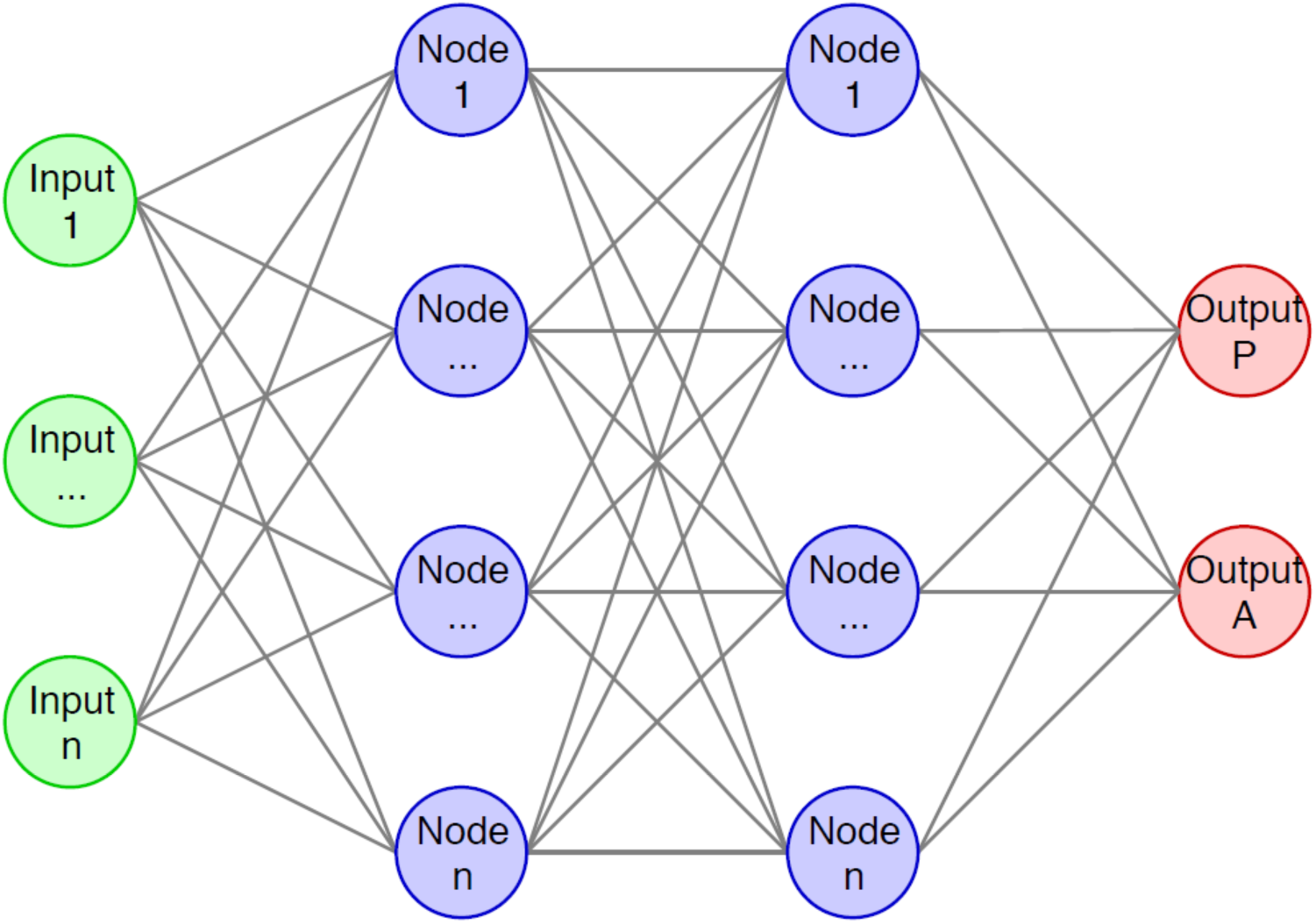
Basic structure of a deep neural network. In green the inputs to the model, in blue the hidden layers and in red the outputs of the model. In this example, output P represents the predicted potential presence of a species while output A represents the predicted potential absence.

An important part of a created model are the hidden layers (represented in blue in Figure 7). A hidden layer can contain different numbers of ‘nodes’ or neurons, and a deep learning model can have two or more hidden layers. In general, shallower models - or models with a small number of hidden layers - are better at generalizing, while deeper models with a larger number of hidden layers are able to model more complex situations (e.g. when using a large number of input variables).

The architecture used in the case study analysis, the default for the package, was relatively shallow: it contained four hidden layers with respectively 250, 200, 150 and 100 nodes (for a visual representation of the default architecture see Figure 8). Each hidden layer was subject to a procedure called ‘dropout’, which turned off a percentage of the layer’s neurons. This prevented overfitting behavior since the model was unable to strengthen the same neural pathways on each training iteration. The dropout percentages were 30%, 50%, 30% and 50% for layers one through four, respectively.

**Figure 8:**
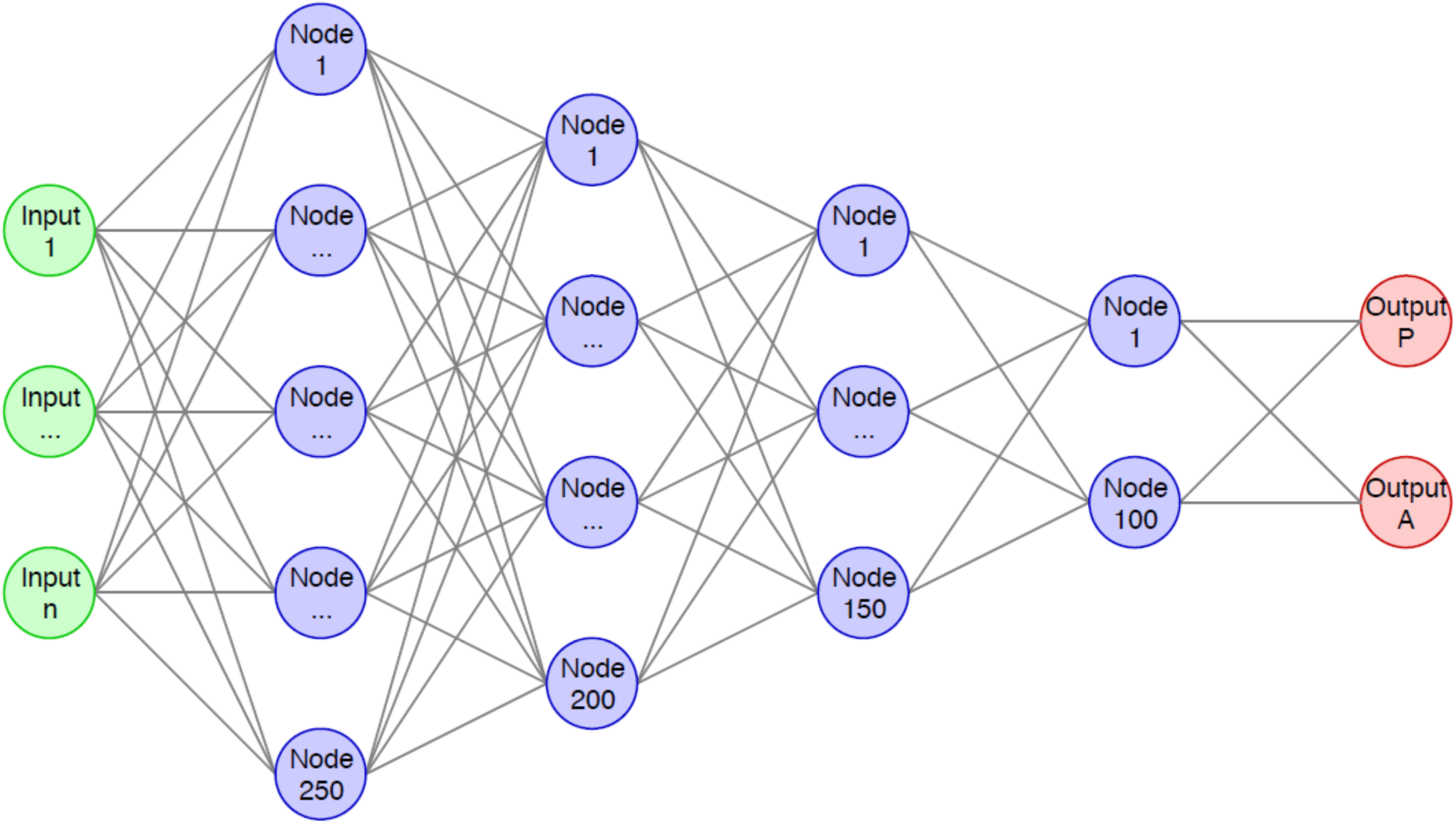
Default model architecture in ‘sdmdl’. In green the inputs to the model, in blue the four hidden layers and in red the output of the model, which in the case of this study is either output P, the predicted potential presence of a species, or output A, the predicted potential absence.

For other cases, the package offers options to change the model architecture by modifying two lists provided in the configuration file. The first list consists of four values by default, which are: 250, 200, 150 and 100, i.e. the number of neurons for each hidden layer. The user is able to change the model architecture by changing the values in the list (e.g. 250, 300, 250, 200) or by adding or removing values to or from the list (e.g. 250, 200, 150). This increases the control the user has over the model, while still remaining user friendly. Additionally, the dropout can also be adjusted using a similar setting, where the values in the list represent the dropout percentage in each layer instead. During model creation a number of model parameters have fixed default values that normally are not exposed to the user. For more technical details on these model parameters see Rademaker et al. [13].

#### DNN training

The training procedure applied here was relatively straightforward, following these steps:

1. A training data handler that handles the input to the model was created.
2. Trained the model for 150 epochs, i.e. full iterations over the training dataset.
3. Evaluated the performance of the model by predicting the presence or absence on a testing dataset.
4. Calculated various performance metrics (test loss, test accuracy, lower confidence interval and upper confidence interval) for later reference.

For more technical details on the training protocol see Rademaker et al. [13].

To allow freedom over the training procedure for other cases, several relevant training parameters are configurable. These parameters are: batch size, i.e. the number of training examples processed by the model at once, 75 by default; number of epochs, i.e. number of training iterations, 150 by default; and random seed, which allows the user to deterministically repeat the creation of a model if an identical value is used for both models, set to 42 by default.

#### Model evaluation

To obtain a viable model, multiple models are created and tested. During this process five different models are created, each initialized with a different set of weights. After their creation, these models are automatically validated to determine which has achieved the best performance. Validating the performance of a deep learning can be done using a number of different performance metrics. The metrics that have been used during this study are Area Under the Curve & Receiver Operating Characteristics curve (AUC & ROC curve) [24]. The value of the ROC curve and AUC score were derived from a number of fundamental metrics of each model, namely the true positive (TP), true negative (TN), false positive (FP) and false negative (FN). A visual representation of these performance metrics can be seen in Figure 9.

**Figure 9:**
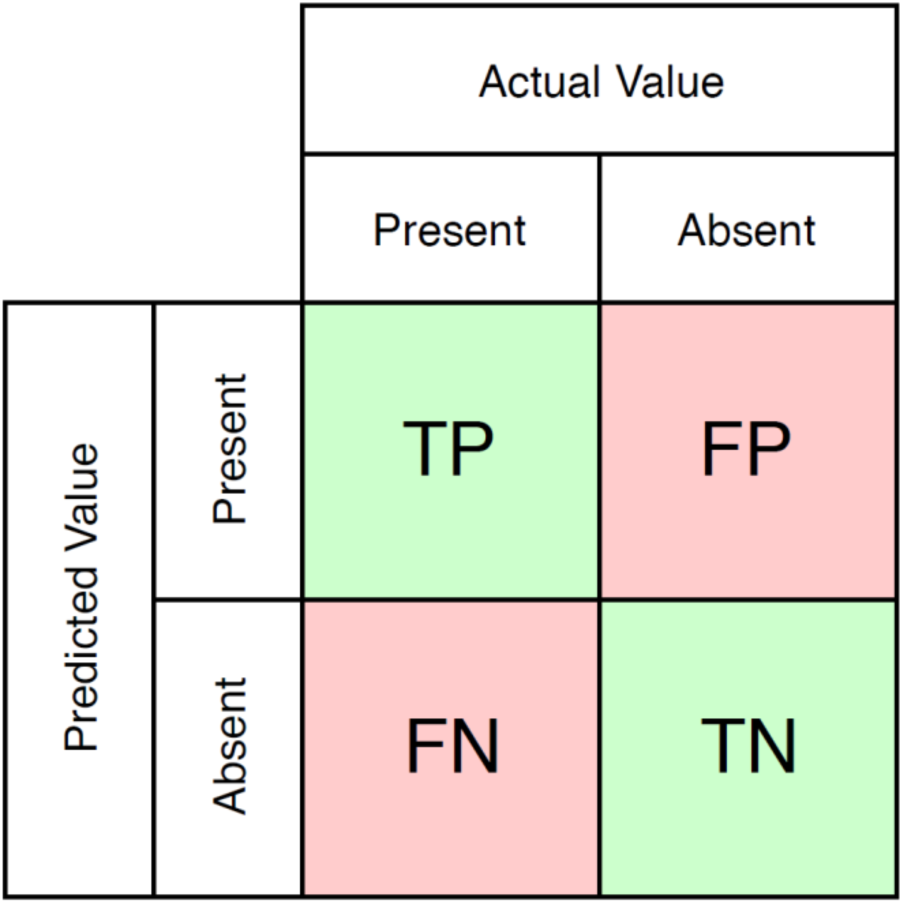
Confusion matrix showing what the true positive (TP), false positive (FP), false negative (FN) and true negative (TN) values represent.

Using these fundamental metrics, the true positive rate (TPR) and false positive rate (FPR) were calculated using the following formulae:

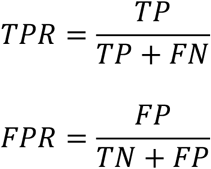

Plotting the TPR versus FPR as a function of the threshold value creates a graph called the ROC curve (for more details see Rademaker et al. [13]). Once this curve is defined the AUC score is calculated by measuring the area under the ROC curve, which is 0.5 if a model is equally likely to predict one class over the other (50% accuracy) and 1 if all predictions are correct (100% accuracy). Out of the five models that were created for each species the one that achieved the highest AUC score was serialized to file.

#### Feature importance

After the single, optimal model was selected based on the model evaluation, the model performed additional computation to determine the impact of each feature (i.e. environmental variable, co-occurrence) used by the model. To achieve this, an approach called shapely values[25] was used. This approach, which originates from game theory, is used to calculate the difference between a model incorporating all variables and a model incorporating all but one variable, thus testing the difference in outcome based on the missing variable. For a more detailed description of the shapely values see Rademaker et al. [13].

#### Distribution prediction

Once the models were trained, the global distribution for a species was predicted. This used a prediction dataset that contained the values of the environmental variables for every terrestrial location on the globe. The model then predicted the presence or absence of a species for each of these locations. Once all the predictions were performed, they were aggregated into a two-dimensional matrix that was then converted into a prediction map and raster layer.

## Results

### Software development

#### Software

During this study a python package called ‘sdmdl’ was created for Python 3.6. This package is open source and available under the MIT license, from the ‘sdmdl’ GitHub repository. This license allows the user to:

1. Use the package commercially.
2. Modify the source code of the package.
3. Distribute modifications of the package.
4. Use the package privately.

Furthermore, it limits any liability and warranty for use of this package. The user is free to use the package as long as they follow these guidelines and correctly cites it. The package was coded using an object oriented programming (OOP) approach, following the PEP-8 style guide for python code [26].

The package contains twelve classes, which can be grouped into three broad groups: Helper, Handler and Main classes. Helper classes perform very specific functions. Handler classes perform more broad processes like managing spatial or occurrence data and model settings. The main class limits the users’ interaction to one class, which only contains a handful of functions needed for performing the entire modeling process. For an overview of each class that is part of the final package and their respective functions see Table 2.

**Table 2:** Class functionality. Showing the class type (helper, handler or main) and a short description of its function.

| Class Name | Type | Function |
| --- | --- | --- |
| Occurrences | Handler | Handles and detects the input and output locations of any occurrence data. |
| GIS | Handler | Handles and detects the input and output locations of any spatial data. |
| Config | Handler | Handles and detects the configuration file and the settings within it. |
| BandStatistics | Helper | Calculates the mean and standard deviation of a raster stack. |
| TrainingData | Helper | Prepares the dataset used for training the models. |
| PredictionData | Helper | Prepares the dataset used for global prediction. |
| Trainer | Helper | Trains the models and performs additional training related functions. |
| Predictor | Helper | Predicts the global habitat suitability and creates a habitat suitability map and raster file. |
| sdmdl | Main | Allows the user to interact with the rest of the packages functionality. Using an interface optimized for ease of use. |

Figure 10 shows how these classes interact with each other. Two types of interactions are visualized, the initiation of an object within another object and one object that serves as the input to another class during its creation (represented in Figure 10 by using an arrow and diamond shape respectively). When executing the main ‘sdmdl’ class its interactions can be grouped chronologically: (1) initiation of an sdmdl object which will create the three handler objects: Config, Occurrences and GIS; (2) data preparation, during which the following objects are created: PresenceMaps, RasterStack, PresencePseudoAbsence, BandStatistics, TrainingData and PredictionData; (3) model training that creates a Trainer object and (4) prediction, where a Predictor object is created.

**Figure 10:**
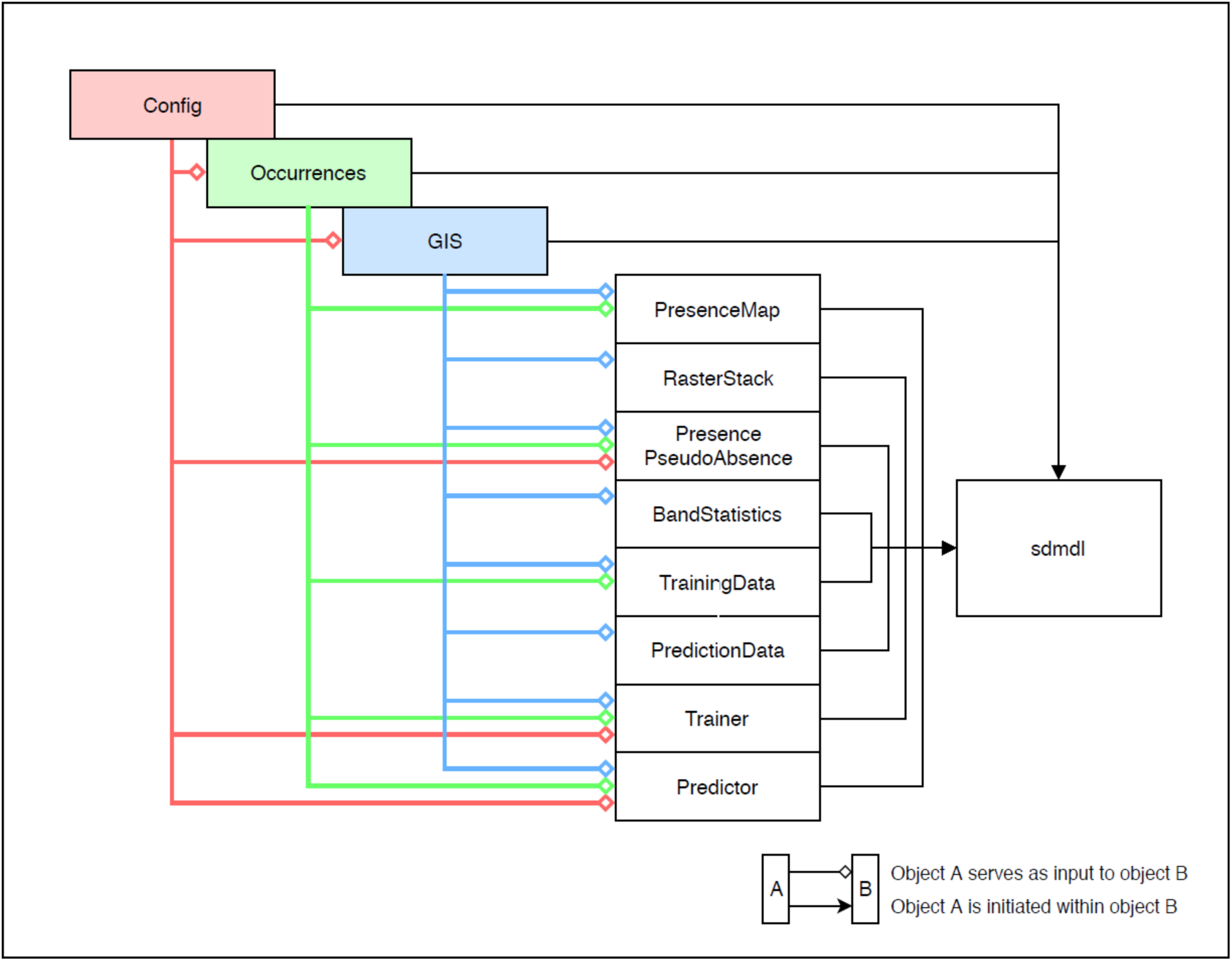
Class interactions within the ‘sdmdl’ package. Showing the handler classes in color, and the helper an main classes in black.

#### Testing

During this study, tests have been performed to ensure the reproducibility of the package. The end result of these testing procedures can be found in Table 3.

**Table 3:** Coverage of unit tests per file from highest to lowest, showing the total coverage per file (in percent), lines of code covered by the tests and lines of code missed by the test. For more details on the package coverage visit https://coveralls.io/github/naturalis/sdmdl?branch=master.

| Coverage | File name | Lines of code covered | Lines of code missed |
| --- | --- | --- | --- |
| 98.51 | sdmdl/gis.py | 66 | 1 |
| 98.46 | sdmdl/data_prep/training_data.py | 64 | 1 |
| 98.33 | sdmdl/data_prep/prediction_data.py | 59 | 1 |
| 96.55 | sdmdl/data_prep/presence_pseudo_absence.py | 56 | 2 |
| 96.55 | sdmdl/data_prep/band_statistics.py | 28 | 1 |
| 96.43 | sdmdl/data_prep/presence_map.py | 27 | 1 |
| 91.67 | sdmdl/data_prep/raster_stack.py | 11 | 1 |
| 91.46 | sdmdl/config.py | 75 | 7 |
| 88.89 | sdmdl/occurrences.py | 40 | 5 |
| 56.21 | sdmdl/trainer.py | 95 | 74 |
| 30 | sdmdl/predictor.py | 27 | 63 |
| 0 | sdmdl_main.py | 0 | 82 |
| **69.63** | **Total** | **548** | **239** |

Most testing performed during this study was relatively straightforward. Tests have generally been performed in two ways. First, if a function or method returns a value, this was tested by saving a visually inspected version of the output to file (most often outputs were numerical arrays) and testing if the values returned by the function or method matched the expected output that was saved to the file. Second, if a function or method saved the output directly to a file already, a similar approach was used which again involved creating a visually inspected copy of the file and matching the file created by the function or method.

It was not possible to test all the code in the package for a number of reasons: (1) any code involving printed feedback to the user was not tested as this would not directly influence the functionality of the package, (2) deep learning code was too complicated to test within the given time due to small disparities between model weights on each run. Another important reason were differences in weight initialization and conditioning between the windows operating system (used on the machine where the tests were created) and the Linux operating system used by the Travis CI service to test the package, and (3) the main class ‘sdmdl_main.py’ was not tested as it mainly uses classes and methods that had already been individually tested.

### ‘sdmdl’ analysis

This section presents the results of applying ‘sdmdl’ to the species occurrences obtained for the case study. Exhaustive discussion of all resulting distribution maps for all species is omitted. Rather, species that have relevant or noteworthy distributions are discussed. The results thus presented fall in the following categories: (1) species with 100 or fewer occurrences, (2) species with 1,000 or fewer occurrences, and (3) species with more than 1,000 occurrences. Once the resulting distribution maps have been discussed a feature importance graph is shown to discuss how these are assessed given the approach presented here. Distribution maps and feature importance graphs for species that are not presented in the results can be found at doi: 10.5281/zenodo.3460718.

#### Species with 100 or fewer occurrences

To illustrate the performance of the package on species with little occurrence data, the predicted potential distribution is discussed for four species with fewer than 100 available occurrences. These species are the ancestors of rye (i.e. *Secale vavilovii*), taro (*Colocasia formosana*), purple yam (*Dioscorea hamiltonii*) and banana/plantain (*Musa balbisiana*). *Secale vavilovii* only had 10 occurrences available. This low number of occurrences has an evident effect on the model, as it is practically unable to learn which locations are suitable or unsuitable to the species (see Figure 11). Instead, the projected suitability is almost global. Moreover, the model has not only predicted worldwide suitability for the species, but has also predicted that this entire distribution is only semi suitable.

**Figure 11:**
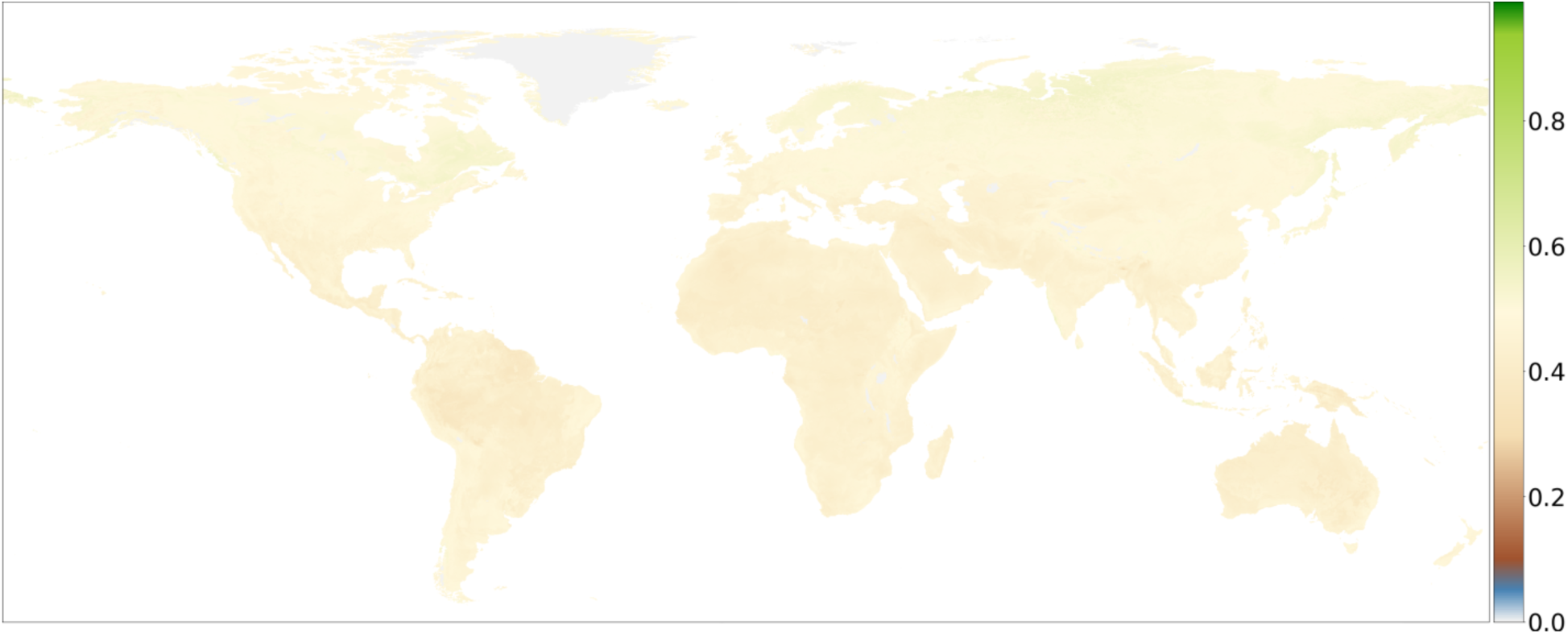
Habitat suitability projection for Secale vavilovii (n_occurrences_=10).

*Colocasia formosana* shows similar results to those of *Secale vavilovii*, likewise showing the inability of the model to learn habitat suitability for a species using a low number of occurrences (see Figure 12). As this species has slightly more occurrences (23), the model is marginally better at deciding which locations are more or less suitable for the species. When compared to the previous case, the distribution map of *Colocasia formosana* shows suitable areas (values close to 1) and unsuitable areas (values of 0.6 or less). Even though the model was better able to distinguish between suitable and unsuitable areas, it predicts a high suitability (suitability = 1) for locations (e.g. Central Africa, Central America and South America) that seem implausible given that it is native to the island of Taiwan [27]. These predictions are most likely side effects of the model being trained with only 23 occurrences, and should not be considered more suitable then areas that were predicted to be mostly suitable (suitability ≈ 0.9).

**Figure 12:**
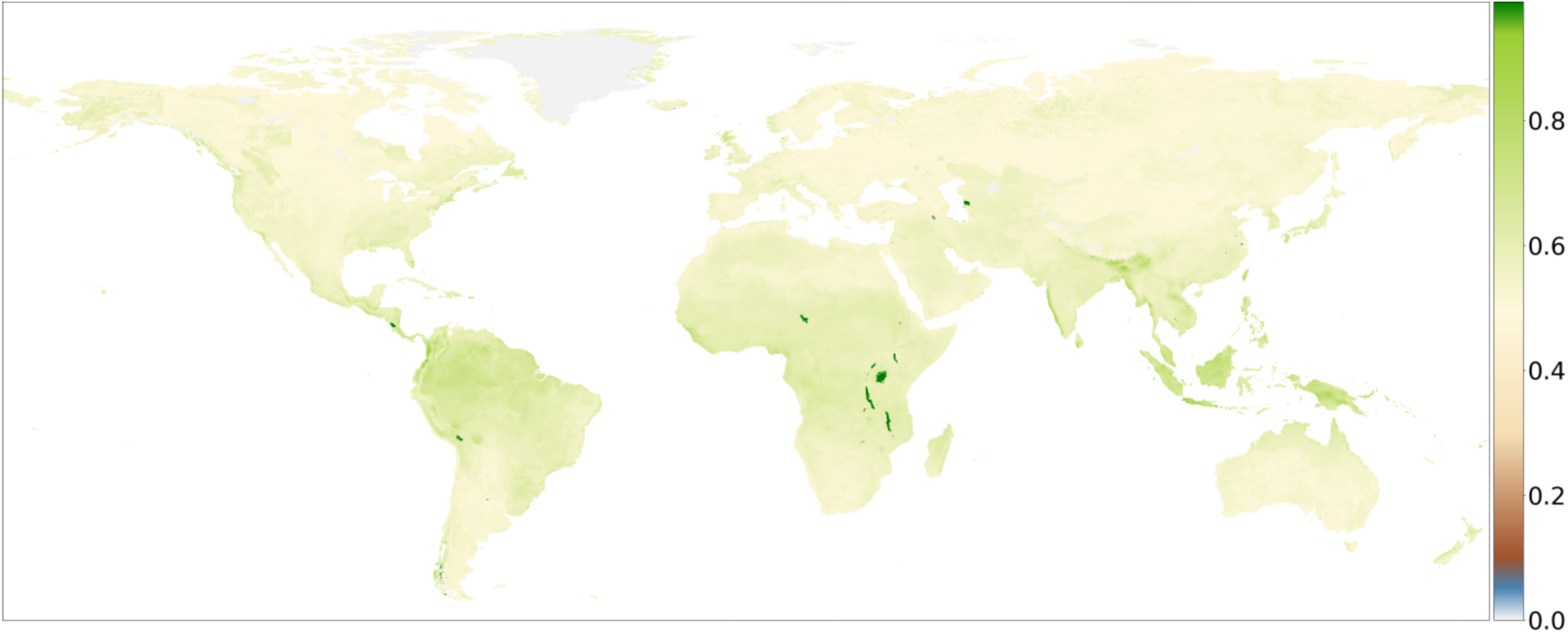
Habitat suitability projection for Colocasia formosana (n_occurrences_=23).

Figure 13 shows the results for the species *Dioscorea hamiltonii*. This map makes it evident that the model is marginally better at distinguishing what makes a habitat suitable, but more importantly, what makes a habitat unsuitable. Even though there is a greater difference between suitable and unsuitable locations, the model still predicts a habitat suitability in locations that should be unsuitable, like Siberia, northern Canada, Greenland and even the most southern extent of South America. This is especially relevant considering that *Dioscorea hamiltonii* is a species with a native range in Southeast Asia [28].

**Figure 13:**
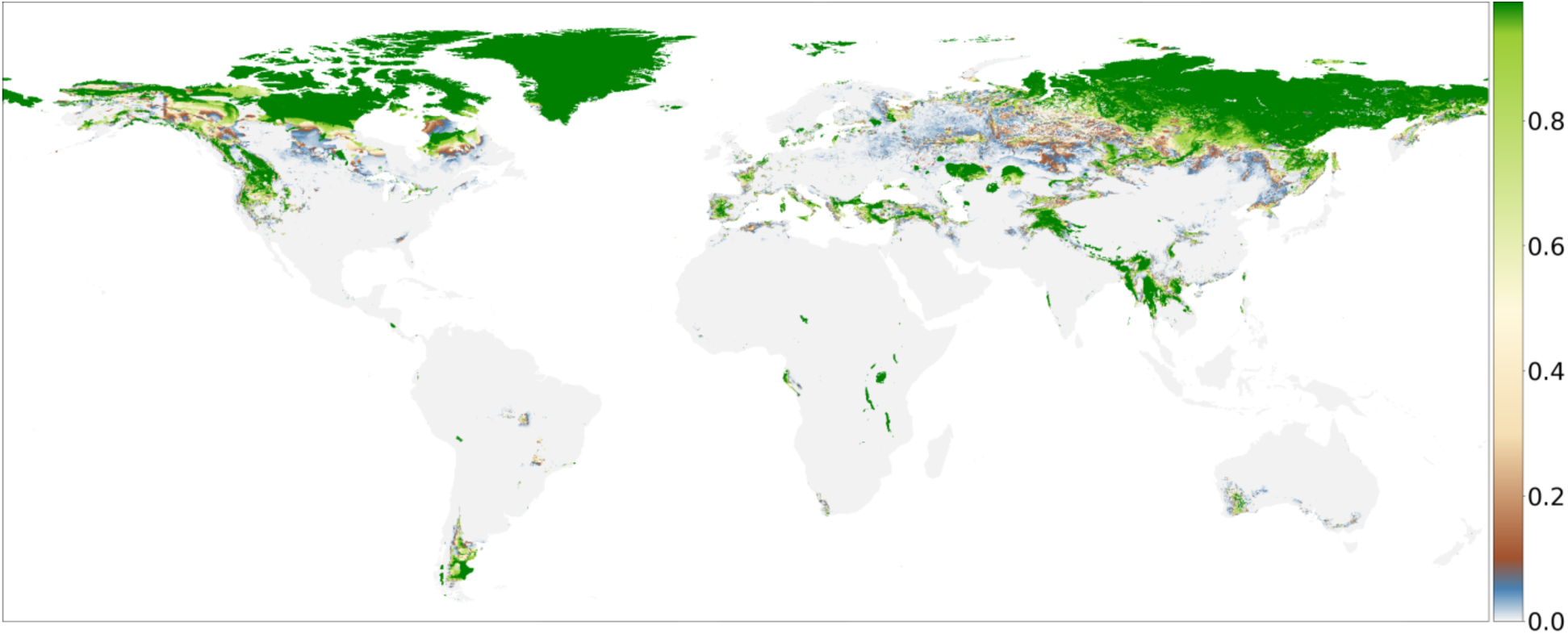
Habitat suitability projection for Dioscorea hamiltonii (n_occurrences_=93).

The last species with 100 occurrences or fewer is *Musa balbisiana*. The suitability projection in Figure 14 shows that the model was mostly able to learn what environmental variables are suitable for the species and which variables made an environment unsuitable. This manifests in the projection mostly covering Southeast Asia, which is where this ancestor originates from [29]. When continents outside of the native range are not considered (e.g. Africa, South and Central America) the model was able to learn what variables make a location suitable for this species. This is especially relevant considering that this species only had 100 occurrences available for training the model.

**Figure 14:**
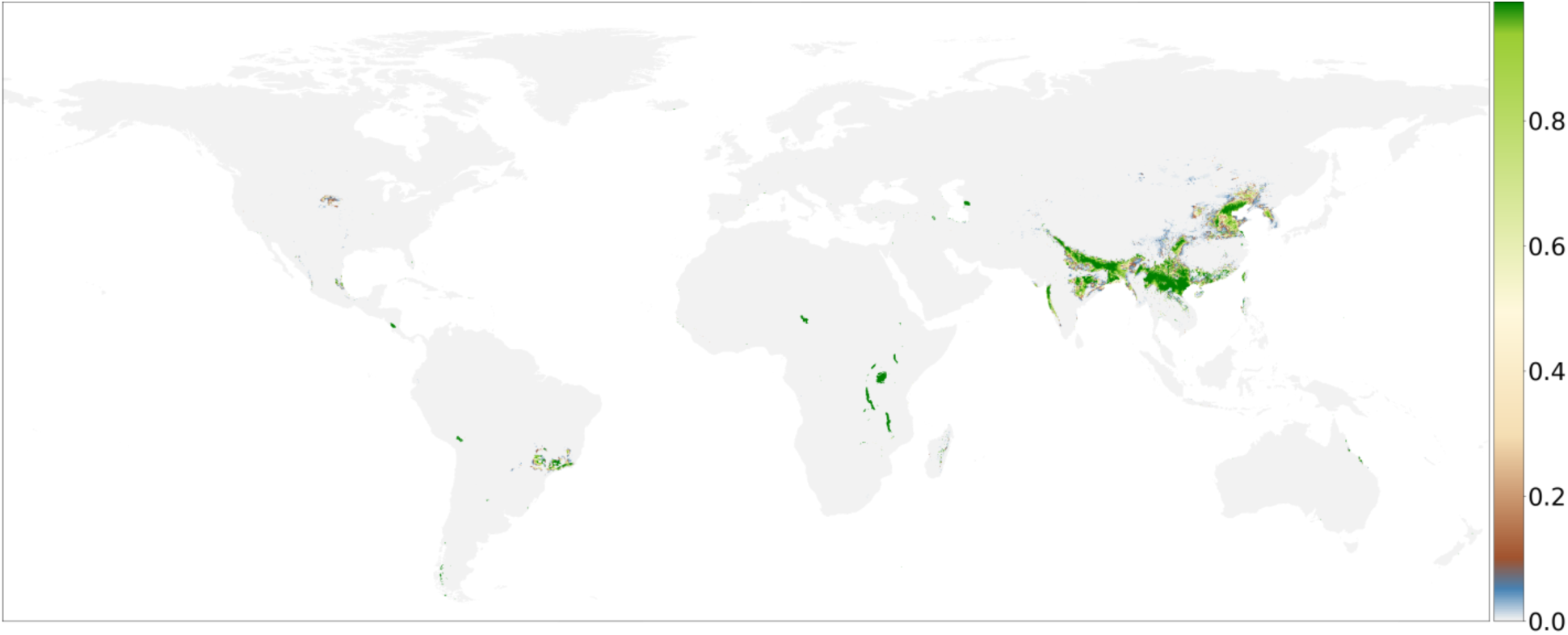
Habitat suitability projection for Musa balbisiana (n_occurrences_=100).

#### Species with 1,000 or fewer occurrences

This section shows the performance of the package on species with more occurrences. To achieve this, the projections for another four species are discussed, which each have up to 1,000 occurrences available. These species are the ancestors of broom corn (i.e. *Sorghum bicolor* subsp. *verticilliflorum*), finger millet (*Eleusine africana*), teosinte, i.e. the ancestor of maize/corn (*Zea mays* subsp. *parviglumis*), and one of the ancestors of common wheat (*Aegilops tauschii*).

Figure 15 shows that the model has over-predicted habitat suitability for *Sorghum bicolor* subsp. *verticilliflorum*. Not only has it over-predicted the spatial range, but also the suitability within that range all around the world. Taking a closer look, Figure 15 reveals that the model has predicted suitability for a range of different climate zones, ranging from the arctic (e.g. North America, Greenland, Scandinavia and Siberia) to the tropics (e.g. the Amazonian basin, India, Southeast Asia and Indonesia). However, the projection does show that the model has been able to learn the suitability for the species on the African continent, which is where this species originates from [30].

**Figure 15:**
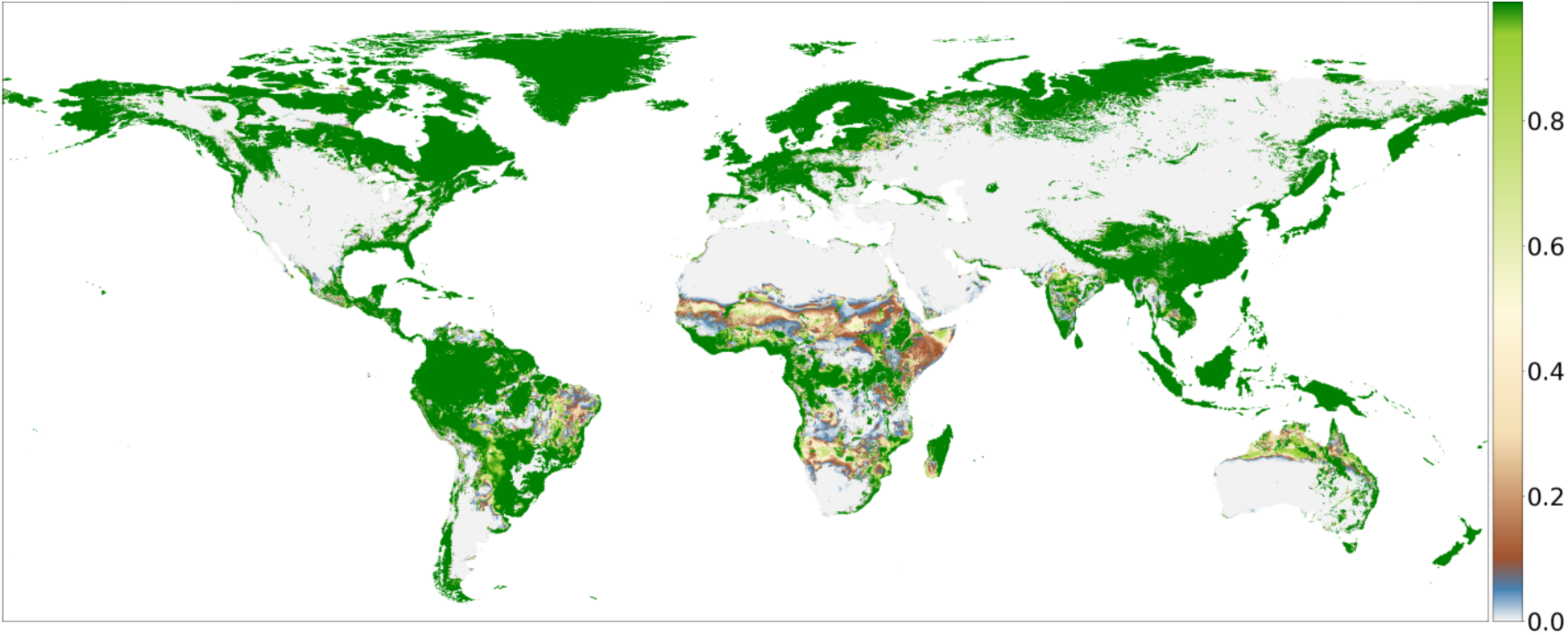
Habitat suitability projection for Sorghum bicolor subsp. verticilliflorum (noccurrences=154).

*Eleusine africana* shows results similar to the previously discussed species (see Figure 16). However, this species shows a more spatially confined projection. The model still projects habitat suitability on every continent, including a wide range of different climate zones, from arctic areas like Iceland and Scandinavia to (sub)tropical areas like the Philippines and Indonesia. When considering the native range of the species, which largely overlaps with sub-Saharan Africa, the predicted suitability more closely matches descriptions of the distribution found in literature, being present in much of the dryer areas in sub-Saharan Africa [31].

**Figure 16:**
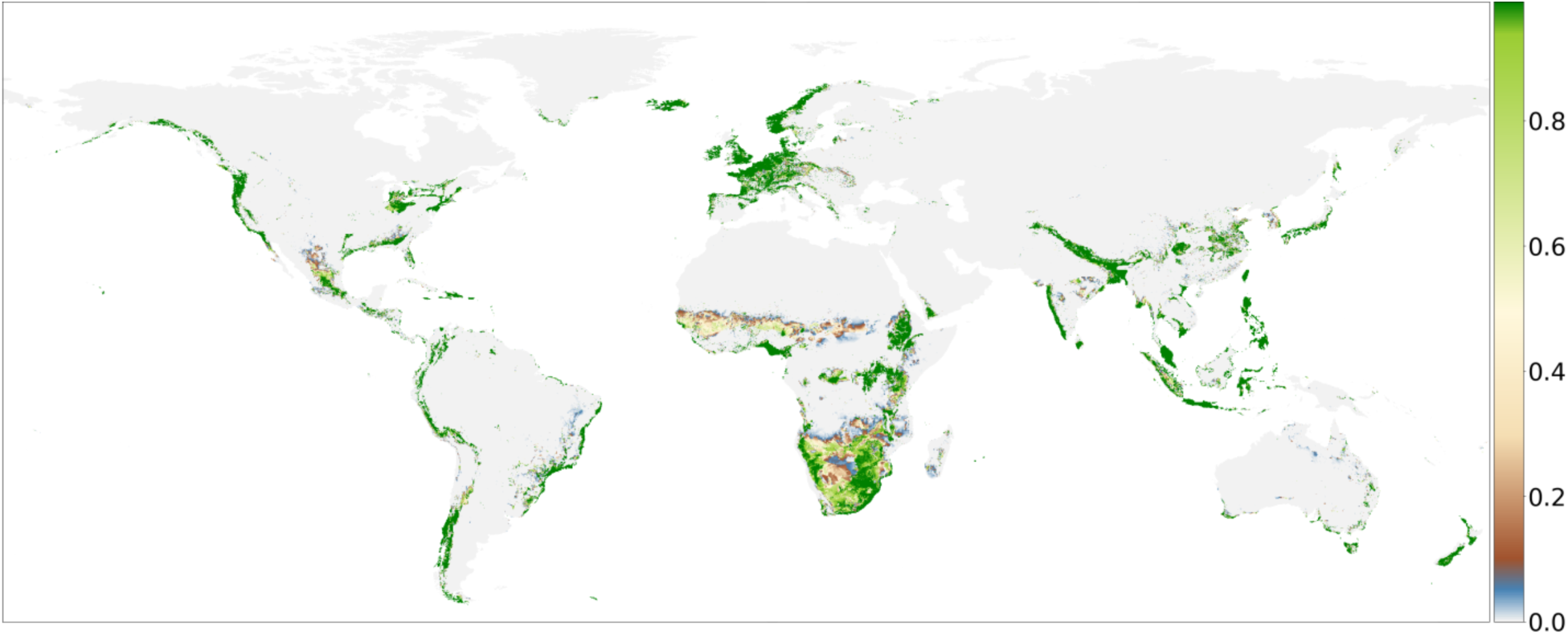
Habitat suitability projection for Eleusine africana (n_occurrences_=357).

*Zea mays* subsp. *parviglumis* shows plausible results. Figure 17 shows that the model was able to learn what locations are suitable for the species, but it also what places are unsuitable. This has led the model to project habitat suitability in line with expectations for the species, while minimizing over-predicting on other continents. As Figure 17 shows, the main location where suitability for this species is projected is in Central America (which closely matches its native distribution) [32], but there are other locations, like Ethiopia, where the model has predicted suitability for this species on a very local scale.

**Figure 17:**
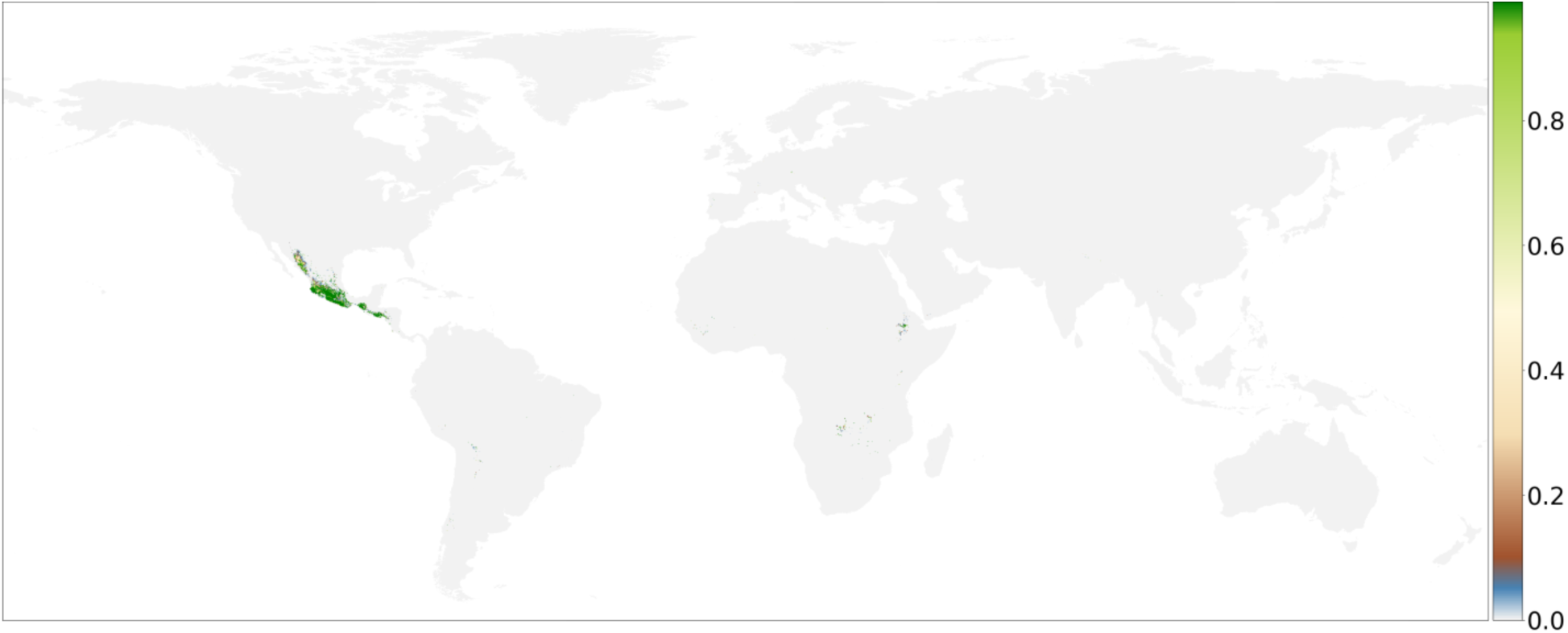
Habitat suitability projection for Zea mays subsp. parviglumis (n_occurrences_=617).

Figure 18 shows the projected suitability for *Aegilops tauschii*. Similarly to the previous discussed species the projection closely matches the distribution described in literature [33]. This means that the model was able to learn to predict suitable locations while minimizing over predicting the species distribution on other continents, or predicting the presence of the species in other climate zones. Apart from the native distribution in the Fertile Crescent the model has also predicted the (local) presence of the species in the north of the African continent and North America.

**Figure 18:**
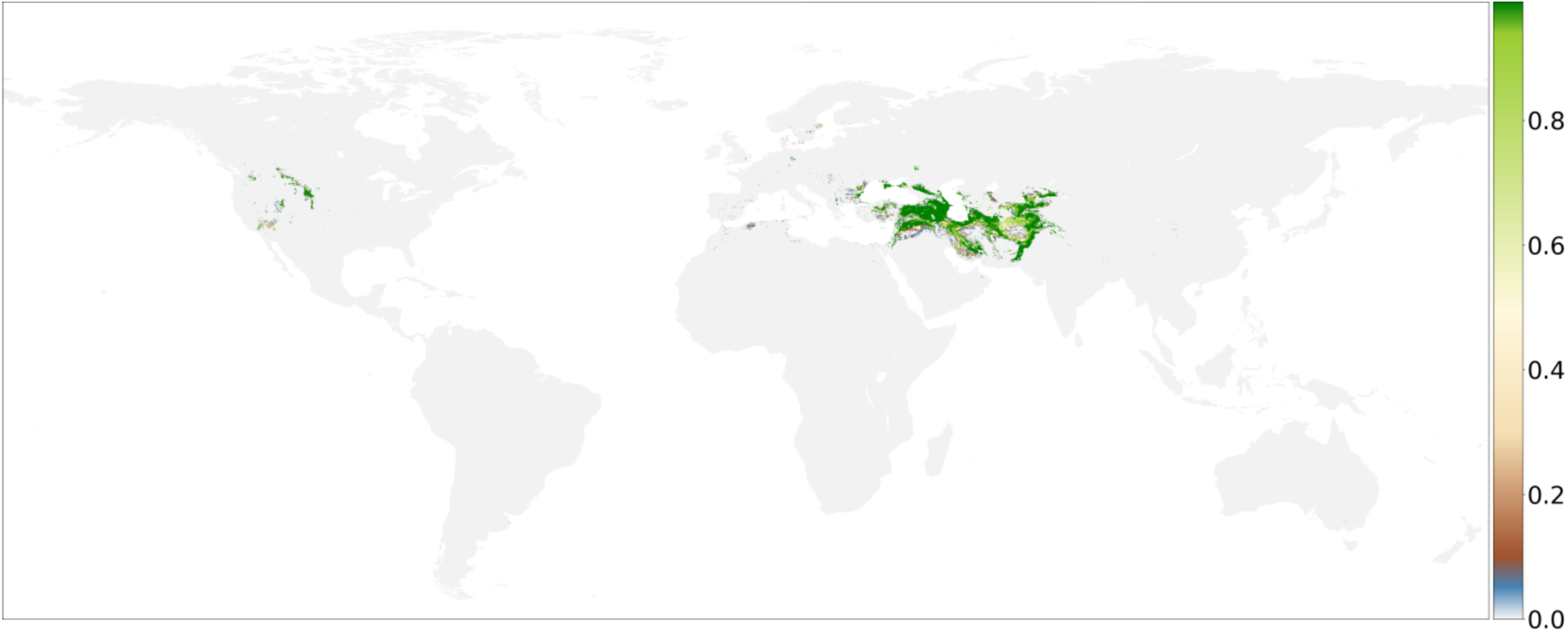
Habitat suitability projection for Aegilops tauschii (n_occurrences_=800).

#### Species with more than 1,000 occurrences

The model performed considerably better when it could be trained using a large number of occurrences and pseudo-absences. This section presents the projected habitat suitability of four exemplar species that have 1,000 occurrences or more available. These species are the domesticated mango (*Mangifera indica*), domesticated rice (*Oryza sativa*), domesticated common wheat (*Triticum aestivum*), and domesticated corn/maize (*Zea mays*). As all of the above-mentioned species are well documented domesticated species with large distribution areas, they are compared to geographical data created by Monfreda et al. [34]. For each species that is discussed in this section, a map is included in Supplementary Figures that shows the fraction of each pixel (between 0 and 1) that is dedicated to growing and harvesting the crop.

Figure 19 shows the projected suitability for mango (*Mangifera indica*). A number of features is evident when looking closer at Figure 19. First, the projection shows close resemblance with the harvested area fraction map of the mango (see Supplementary Figures I). Most predictions are found within a similar latitudinal range, with an overlap between the predicted distribution and harvested area fraction in central and South America, Africa, India, Southeast Asia, Indonesia and Australia. However, when comparing these maps, the model has slightly under predicted the presence of the mango in parts of India, Pakistan and the east coast of China. Finally, it is noteworthy that with a higher number of occurrences the ability of the model to learn and project suitability increases.

**Figure 19:**
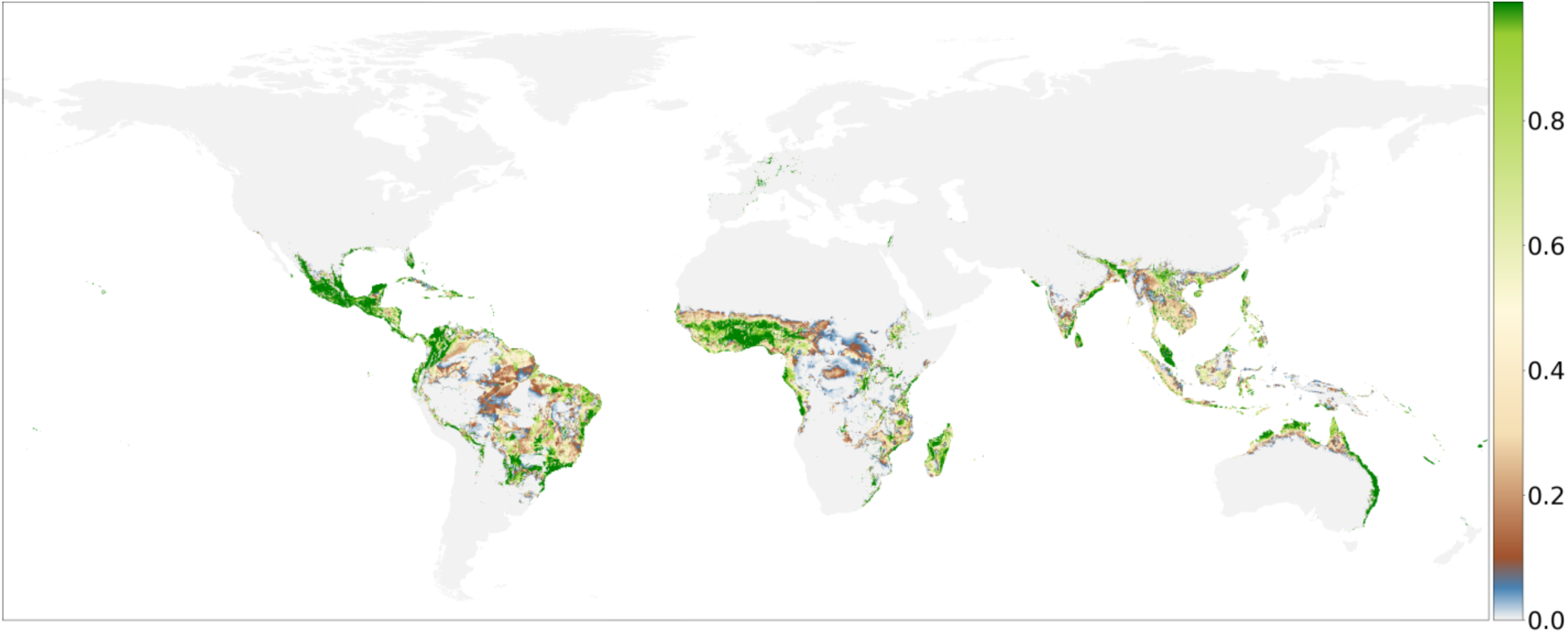
Habitat suitability projection for Mangifera indica (n_occurrences_=2,996).

The projected suitability for rice (*Oryza sativa*) covers a wide range of locations (see Figure 20). Most of the locations that were predicted to have high suitability correspond to the harvested area fraction map in Supplementary Figures II, like Southeast Asia, India, West Africa and Central America. The highest suitability is projected close or slightly north of the equator, with some minor exceptions like Norway where low suitability was predicted. The location with the highest suitability (Southeast Asia) also seems plausible as this is the area where the ancestor of rice (*Oryza rufipogon*) originated.

**Figure 20:**
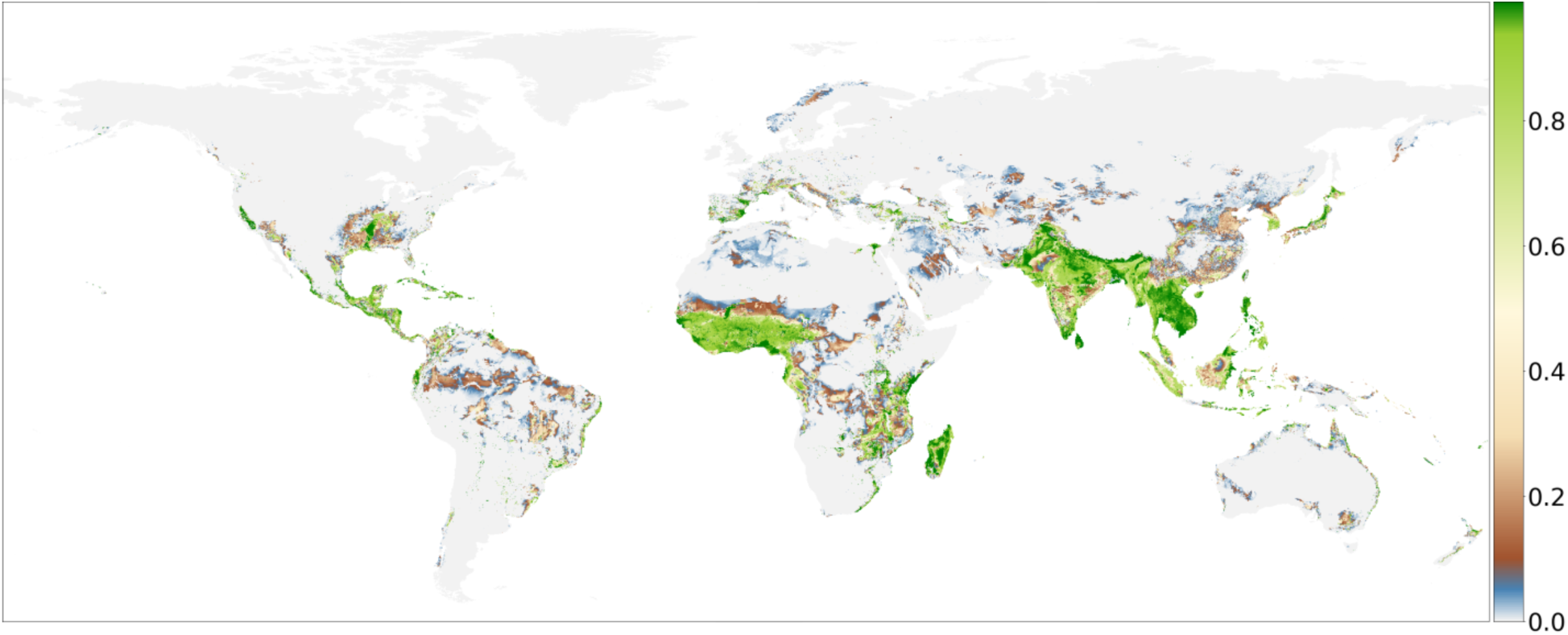
Habitat suitability projection for Oryza sativa (n_occurrences_=7,145).

Common wheat (*Triticum aestivum*) is one of the world’s most consumed crops and as such has a large worldwide distribution. This can be seen in Figure 21, which shows the suitability projected by the model. This figure shows that the species is primarily projected across the northern hemisphere. This is confirmed by the harvested area map in Supplementary Figure III. Locations with high suitability are mainly projected in Europe, with additional projections in central America, the Fertile Crescent, the north of China and Australia. However, some of these locations, like Central America and northern China, seem to be somewhat over predicted, while other areas like North America, eastern china, South America and South Africa seem to be under predicted when compared to the harvested area fraction map. The high suitability projection in the Fertile Crescent is also noteworthy as this is the native distribution of multiple ancestor species of common wheat.

**Figure 21:**
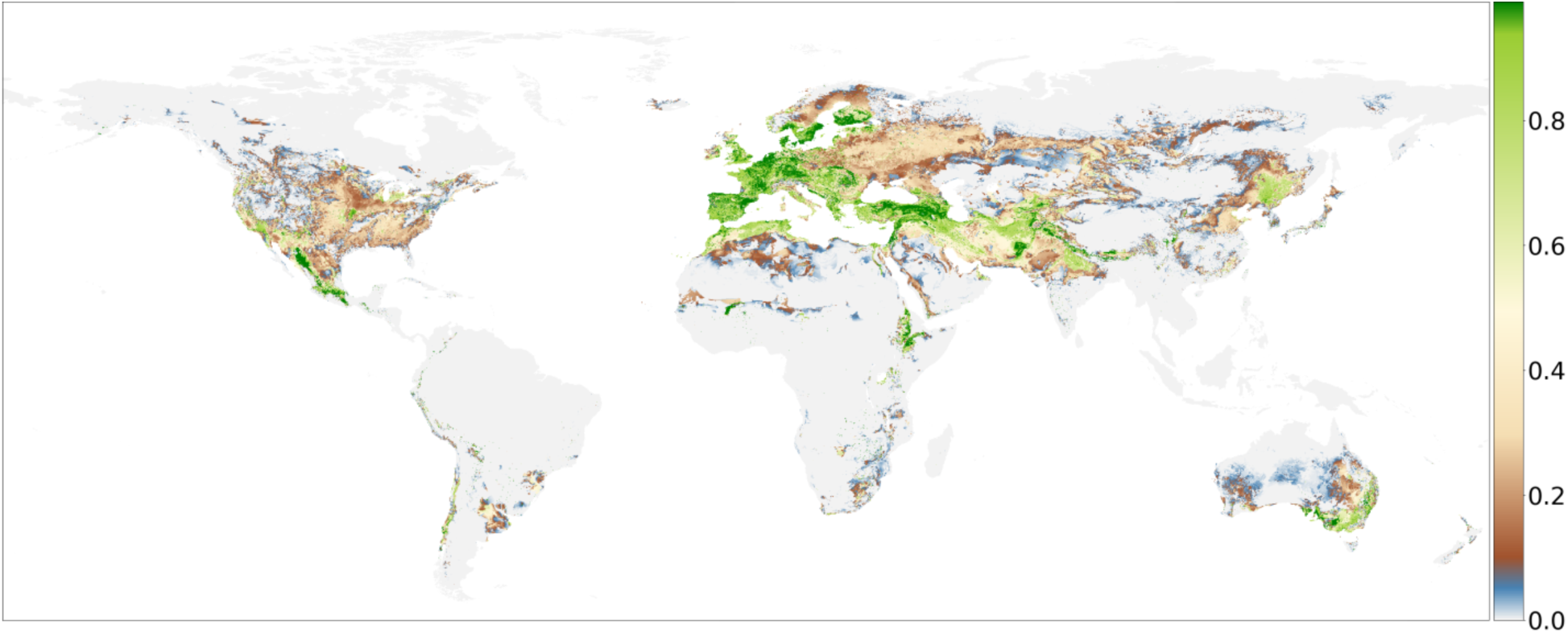
Habitat suitability projection for Triticum aestivum (n_occurrences_=11,572).

The projected suitability for the final species, *Zea mays*, is shown in Figure 22. The model predicted that suitable areas can be found in both the northern and southern hemisphere; however, it is largely missing from the eastern hemisphere. As corn is a very common crop it is noticeably missing from a number of locations when compared to the harvested area fraction map in Supplementary Figure IV. Locations where the model under predicted suitability include North America, northeastern China, South Africa and India. It is, however, noteworthy that the location with the highest suitability is Central America, where the ancestor of the domesticated corn plant originates from.

**Figure 22:**
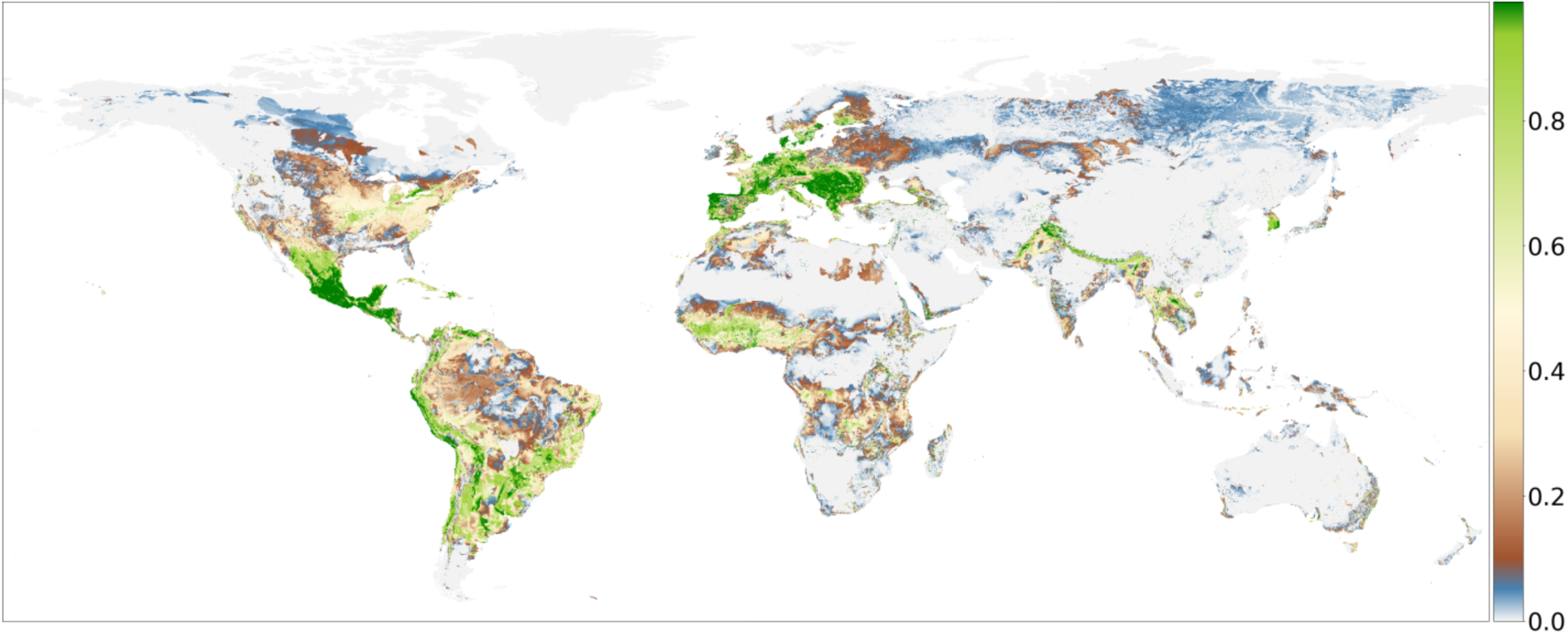
Habitat suitability projection for Zea mays (n_occurrences_=15,495).

#### Feature importance

The importance each variable has in the model can be tested individually by using the shapely values [25]. However, detailed interpretation of the ecological implications of each individual species and variable is omitted; feature importance is discussed by looking at the importance of presence instead. An example of a feature importance graph is given in Figure 23.

**Figure 23:**
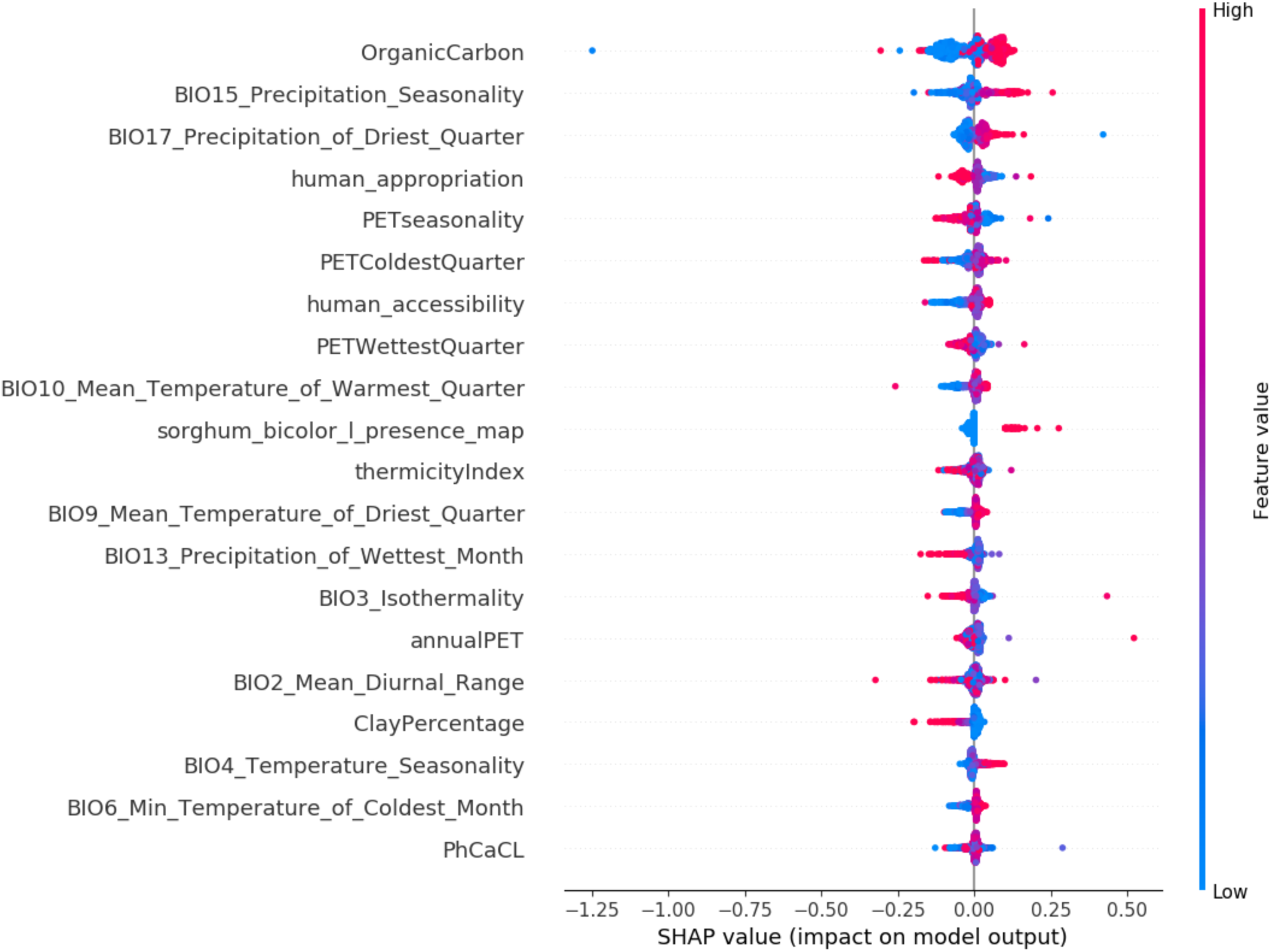
Feature importance graph for Eragrostis pilosa. Showing 20 variables used for the prediction of presence, in order of importance. For each variable the figure shows how the final prediction is influenced by the value of the feature/variable.

In cases where the ancestor and domesticated species of a crop have an overlap in distribution, and assuming that both species have similar ecological niches, it could be expected that the presence of the domesticated species functions as an important variable for predicting habitat suitability for the ancestor. Table 4 shows that in nine out of the total 29 ancestor species the presence of their domesticated counterpart is such an important variable in the prediction of suitability for the ancestor. Conversely, for 20 out of 29 species the presence of its corresponding crop species does not function as a predictor. This high number can partially stem from a lack of occurrences for these species, as sufficient overlap needs to be present for this to be an important variable. This can be seen by looking at number of occurrences for each of the species where the presence of the crop is an important variable. This reveals that most of these species have at least 500 or more occurrences. Interestingly, for some species the presence of their related crop species does not factor into their habitat suitability projection; rather, the presence of another ancestor of the same crop is used (e.g. *Triticum urartu* uses the presence of *Triticum turgidum* subsp. *Dicoccoides*, both ancestors of common wheat). In other cases, the presence of closely related species or species that have originated in the same geographical region are used (e.g. *Aegilops speltoides*, ancestor of common wheat, uses the presence of *Hordeum spontaneum*, ancestor of rye; and *Eragrostis pilosa*, ancestor of teff, uses the presence of *Sorghum bicolor*, broom corn).

**Table 4:** Importance of crop occurrence in habitat suitability projections for ancestor species. Shows for each species (1) crop species (2) scientific name of its ancestor (3) number of occurrences per ancestor and (4) whether the crop species is an important predictor for habitat suitability projection of its ancestor, and if so how important is the variable (value in parentheses corresponds to the importance of the variable, where 1 = most important).

| Crop species | Scientific name of ancestor | n<sub>occurrences</sub> | Crop presence importance |
| --- | --- | --- | --- |
| Teff | *Eragrostis pilosa* | 3639 | (>20) |
| Rice | *Oryza rufipogon* | 3144 | 11 |
| Barley | *Hordeum spontaneum* | 2720 | 16 |
| Sweet potato | *Ipomoea trifida* | 904 | 11 |
| Common wheat [1] | *Aegilops tauschii* | 800 | 9 |
| Common wheat [2] | *Triticum turgidum* subsp. *dicoccoides* | 743 | (>20) |
| Maize [1] | *Zea mays* subsp. *parviglumis* | 617 | 1 |
| Maize [2] | *Zea mays* subsp. *mexicana* | 604 | 4 |
| White yam [1] | *Dioscorea cayennensis* | 474 | 2 |
| Common wheat [3] | *Aegilops speltoides* | 398 | (>20) |
| Cassava | *Manihot esculenta* subsp. *flabellifolia* | 364 | 1 |
| Finger millet | *Eleusine africana* | 357 | (>20) |
| Cushaw pumpkin | *Cucurbita argyrosperma* subsp. *sororia* | 350 | 1 |
| White yam [2] | *Dioscorea praehensilis* | 307 | (>20) |
| Common wheat [4] | *Triticum urartu* | 278 | (>20) |
| Potato | *Solanum bukasovii* | 181 | (>20) |
| Plantain [1] | *Musa acuminata* | 181 | (>20) |
| Yellow yam | *Dioscorea minutiflora* | 167 | (>20) |
| Broom-corn | *Sorghum bicolor* subsp. *verticilliflorum* | 154 | (>20) |
| White yam [3] | *Dioscorea abyssinica* | 135 | (>20) |
| Plantain [2] | *Musa balbisiana* | 100 | (>20) |
| Purple yam [1] | *Dioscorea hamiltonii* | 95 | (>20) |
| Purple yam [2] | *Dioscorea persimilis* | 75 | (>20) |
| Squash | *Cucurbita maxima* subsp. *andreana* | 28 | (>20) |
| Taro | *Colocasia formosana* | 23 | (>20) |
| Rye | *Secale vavilovii* | 10 | (>20) |
| Breadfruit | *Artocarpus camansi* | 9 | (>20) |
| Mango | *Mangifera sylvatica* | 9 | (>20) |
| Dates | *Phoenix sylvestris* | 6 | (>20) |

## Discussion

This paper outlines a generalized approach, presenting its results with reference to functionality of a software package that was a major outcome of this research. The functionality is assessed in terms of the predicted habitat suitability for the species in the case study. However, due to the nature of the case study, in which species were included with relatively low numbers of occurrences, it was not possible to obtain external data sources that could be used to validate the results and test how accurate they are when compared to other data sources. This means that this report is unable to provide metrics on the exact performance of the created models. Nevertheless, the results show some of the general patterns that can be expected in the study of the biodiversity of domestication. Namely, that the process of domestication would suggest that crop species have been selected to survive in a wider variety of habitats. Thus domesticated species should have a wider distribution and are able to thrive in a wider range of niches then their ancestor. This pattern can be seen in a number of predicted crop species including: maize, rice, barley, common wheat, potato, sweet potato and peanut. The most plausible evidence for this patterns is found in species with more than about 500 distinct occurrence points available, as was also found by Rademaker et al. [13]. Another pattern that was observed is related to spatial autocorrelation, more precisely the use of presence maps from close genetic relatives for prediction. In species with a small number of occurrences (e.g. *Manihot esculenta* subsp. *flabellifolia*, the ancestor of cassava) this has been observed to drastically affect the predicted output of the model, essentially creating a model that almost exclusively predicts a high suitability in locations where its genetic relative is present.

The random pseudo-absence sampling procedure used during this study has had an influence on the reliability of the final results, especially for species with little occurrences. This can be largely attributed to the way absences are sampled. As the model is trained on both presence and absence of a species, sampling random locations for absences can have a drastic influence on the final output. For example, if two identical models are created for the same species, which use two different sets of randomly sampled absences, the results can show drastic changes between them. This is because the model learns what locations are unsuitable to a species based on the environmental variables of the randomly sampled absence locations. During one run locations that are potentially suitable to the species may be sampled, conditioning the model to under predict the suitability for the species. On the other hand, the model may only sample locations that are unsuitable for the species. In this case, the opposite will happen and the model will project a considerably larger distribution. This has been observed for some crop species throughout the development of the package. And is a relevant especially for species with little occurrences, as they have a smaller number of pseudo absences.

During the model training the AUC performance metric was used. This can also have an influence on the final performance of the model as the AUC score relies on the distribution between true positives, true negatives, false positives and false negatives. However, as these metrics rely on proper ground truth labels this can become a problem. Since at least half of the data used to train and validate the model is randomly sampled, there is no guarantee that locations that are randomly sampled are not in fact suitable for the species, or that the species might, in fact, be present yet unrecorded. The ambiguity as to whether the species is actually absent, or simply not documented in that location, can affect the final performance metrics in such a way that they become unreliable. This is especially relevant for species with a low number of occurrences, as this can mean that locations that are directly adjacent to occurrences can still be selected as absence locations, while the chance is relatively high that the species is present here yet undocumented. Nonetheless, it is worth to mention that this does not invalidate the model and its performance, but has the potential to provide a slight distortion in performance metrics.

Some functionalities should be considered for implementation in the ‘sdmdl’ package. The most urgent feature is the implementation of functionality regarding the input of environmental raster layers. In the current state the package is unable to use provided raster layers unless they have a specific spatial extent and a resolution (of 5 arc minutes). These requirements can prevent people from using this package, especially in cases in which the user has no experience working with raster layers or other spatial data formats. Other less urgent features are discussed and documented within the code on the GitHub repository.

## Conclusions

This paper presents a proof of concept of user friendly SDM with deep learning and gives insight into how the model performs on the ‘crop and ancestor’ case study. This case study is used to illustrate under what circumstances the model is able to perform as expected while also discussing the model’s potential weak points, which could lead to producing biased, or incorrect results.

The ‘sdmdl’ approach is easy to use by researchers with little experience with programming and deep learning; by using four simple commands the entire procedure can be performed. Furthermore, the model offers a variety of parameters that give the user extensive freedom to change the behavior of the model. These parameters are accessible via an editable configuration file, and allow the user to change settings regarding the species and the environmental variables included in the analysis, the architecture of the model, and several more (see documentation on the GitHub repository for more details on all the configuration parameters at https://sdmdl.readthedocs.io).

This study demonstrates that the methodology used is applicable to other groups of species as the default parameters used for performing the case study are the parameters used by Rademaker et al. [13] to predict the potential distribution of ungulates (hooved mammals). In conclusion, this study has succeeded in using the deep learning methodology devised by Rademaker et al. [13] and adapting it into a package that offers an intuitive interface and considerable freedom for the user to change how the model performs and behaves. However, in its current state, ‘sdmdl’ still needs a number of features that are crucial for it to be broadly applicable. These features will be discussed in the following section.

## Future work

The approach used to sample random absences has been previously reviewed in the discussion section. However, a relatively simple approach could be used to improve this methodology. As previously mentioned the sampled locations can have a profound impact on the predicted potential distribution. Sampling locations close to occurrence points can condition the model to predict suitable locations as unsuitable. One way to prevent this is using a geodesic buffer around occurrences where no pseudo-absences can be sampled. The size of this buffer should be relatively small, to prevent the model from over fitting, but also should not be so small that it might include locations that are likely suitable to the species.

Another way to improve the model would be to find a more suitable performance metric to the nature of the problem, i.e. a metric that does not directly rely on the ground truth labels of the pseudo-absences. However, this could prove a difficult implementation as most performance metrics rely on the ground truth in some way. Some alternatives for deep learning classification performance metrics are precision-recall, accuracy and log-loss [35].

One solution to counteract the shortcomings of the pseudo-absence sampling and AUC-ROC metric would be to use an approach similar to Monte Carlo simulations [36]. This would be especially useful on species with lesser availability of occurrence data. As pseudo-absences have an inherent uncertainty, using multiple models a number of different ‘scenarios’ can be simulated, each model trained using a different set of pseudo-absences. By repeating the analysis multiple times, and finally concluding by taking the mean or median value for each predicted location, the resulting map should be more accurate and less susceptible to the influence of the random sampling of pseudo absences.

## Supplementary materials

**Supplementary Table.**
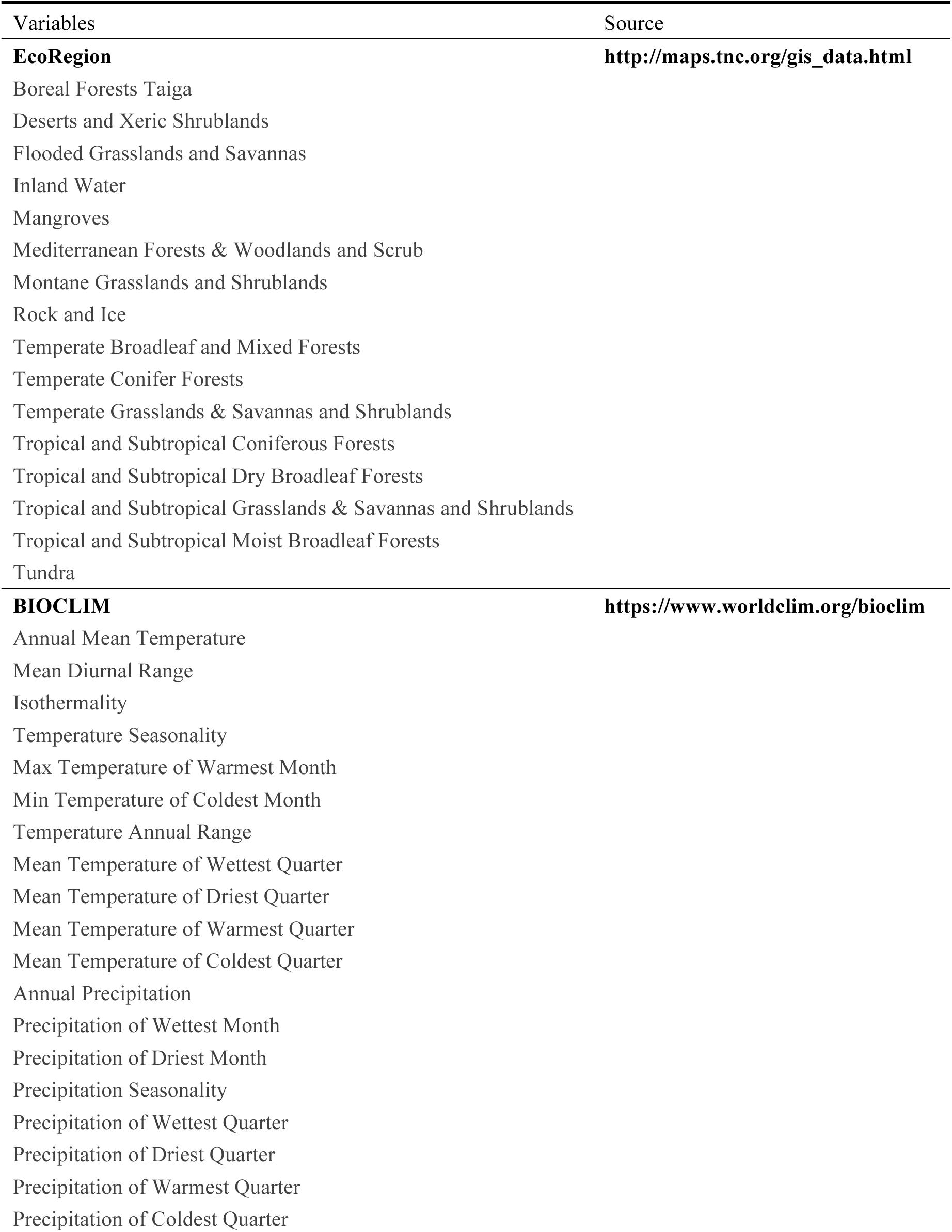
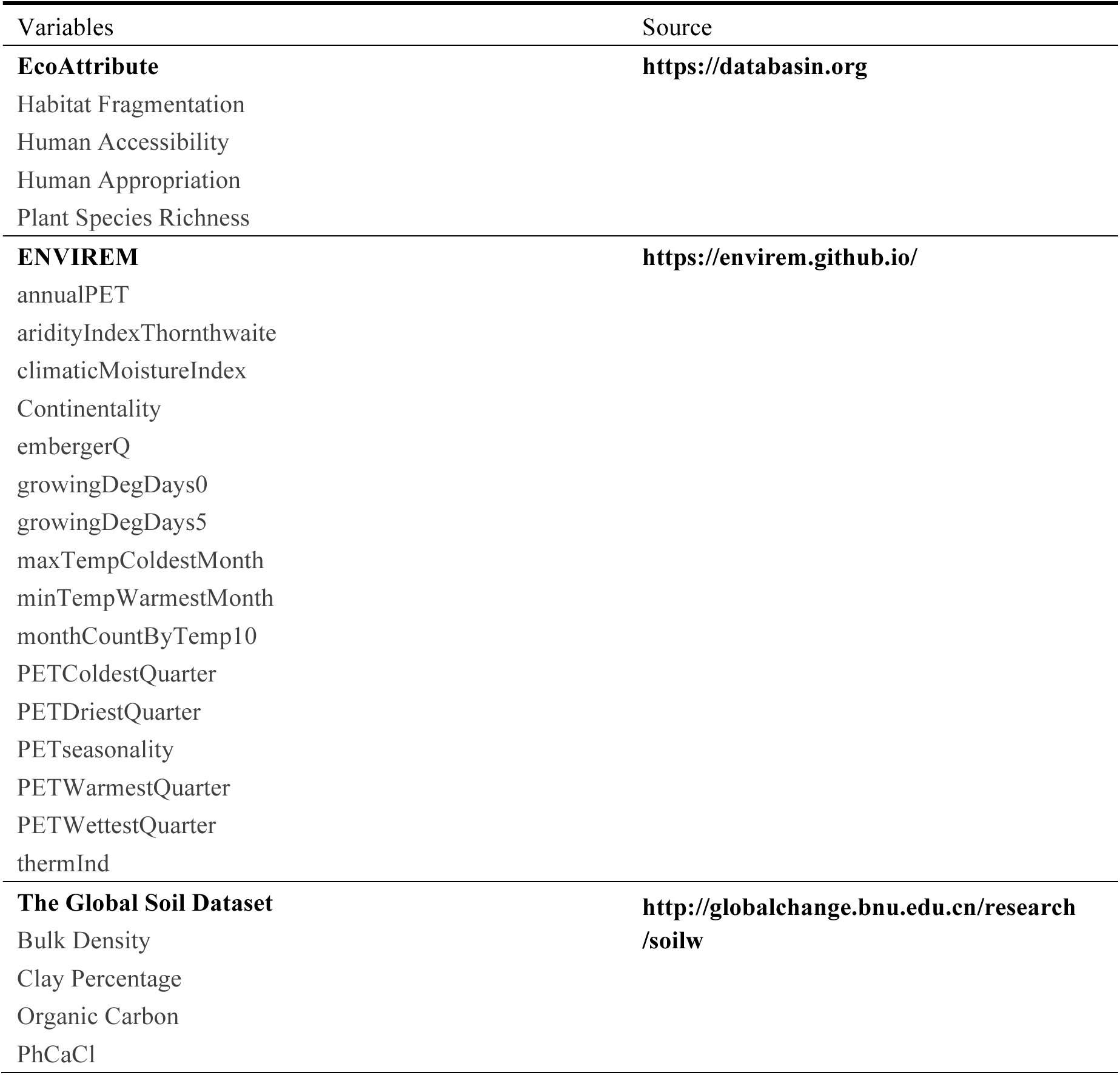
Environmental variables

| Variables | Source |
| --- | --- |
| **EcoRegion** | **http://maps.tnc.org/gis_data.html** |
| Boreal Forests Taiga |  |
| Deserts and Xeric Shrublands |  |
| Flooded Grasslands and Savannas |  |
| Inland Water |  |
| Mangroves |  |
| Mediterranean Forests & Woodlands and Scrub |  |
| Montane Grasslands and Shrublands |  |
| Rock and Ice |  |
| Temperate Broadleaf and Mixed Forests |  |
| Temperate Conifer Forests |  |
| Temperate Grasslands & Savannas and Shrublands |  |
| Tropical and Subtropical Coniferous Forests |  |
| Tropical and Subtropical Dry Broadleaf Forests |  |
| Tropical and Subtropical Grasslands & Savannas and Shrublands |  |
| Tropical and Subtropical Moist Broadleaf Forests |  |
| Tundra |  |
| **BIOCLIM** | **https://www.worldclim.org/bioclim** |
| Annual Mean Temperature |  |
| Mean Diurnal Range |  |
| Isothermality |  |
| Temperature Seasonality |  |
| Max Temperature of Warmest Month |  |
| Min Temperature of Coldest Month |  |
| Temperature Annual Range |  |
| Mean Temperature of Wettest Quarter |  |
| Mean Temperature of Driest Quarter |  |
| Mean Temperature of Warmest Quarter |  |
| Mean Temperature of Coldest Quarter |  |
| Annual Precipitation |  |
| Precipitation of Wettest Month |  |
| Precipitation of Driest Month |  |
| Precipitation Seasonality |  |
| Precipitation of Wettest Quarter |  |
| Precipitation of Driest Quarter |  |
| Precipitation of Warmest Quarter |  |
| Precipitation of Coldest Quarter |  |
| **EcoAttribute** | **https://databasin.org** |
| Habitat Fragmentation |  |
| Human Accessibility |  |
| Human Appropriation |  |
| Plant Species Richness |  |
| **ENVIREM** | **https://envirem.github.io/** |
| annualPET |  |
| aridityIndexThornthwaite |  |
| climaticMoistureIndex |  |
| Continentality |  |
| embergerQ |  |
| growingDegDays0 |  |
| growingDegDays5 |  |
| maxTempColdestMonth |  |
| minTempWarmestMonth |  |
| monthCountByTemp10 |  |
| PETColdestQuarter |  |
| PETDriestQuarter |  |
| PETseasonality |  |
| PETWarmestQuarter |  |
| PETWettestQuarter |  |
| thermInd |  |
| **The Global Soil Dataset** | **http://globalchange.bnu.edu.cn/research/soilw** |
| Bulk Density |  |
| Clay Percentage |  |
| Organic Carbon |  |
| PhCaCl |  |

## Supplementary Figures – Harvested Area Fraction for Domesticated Crop Species

**Supplementary Figure I:**
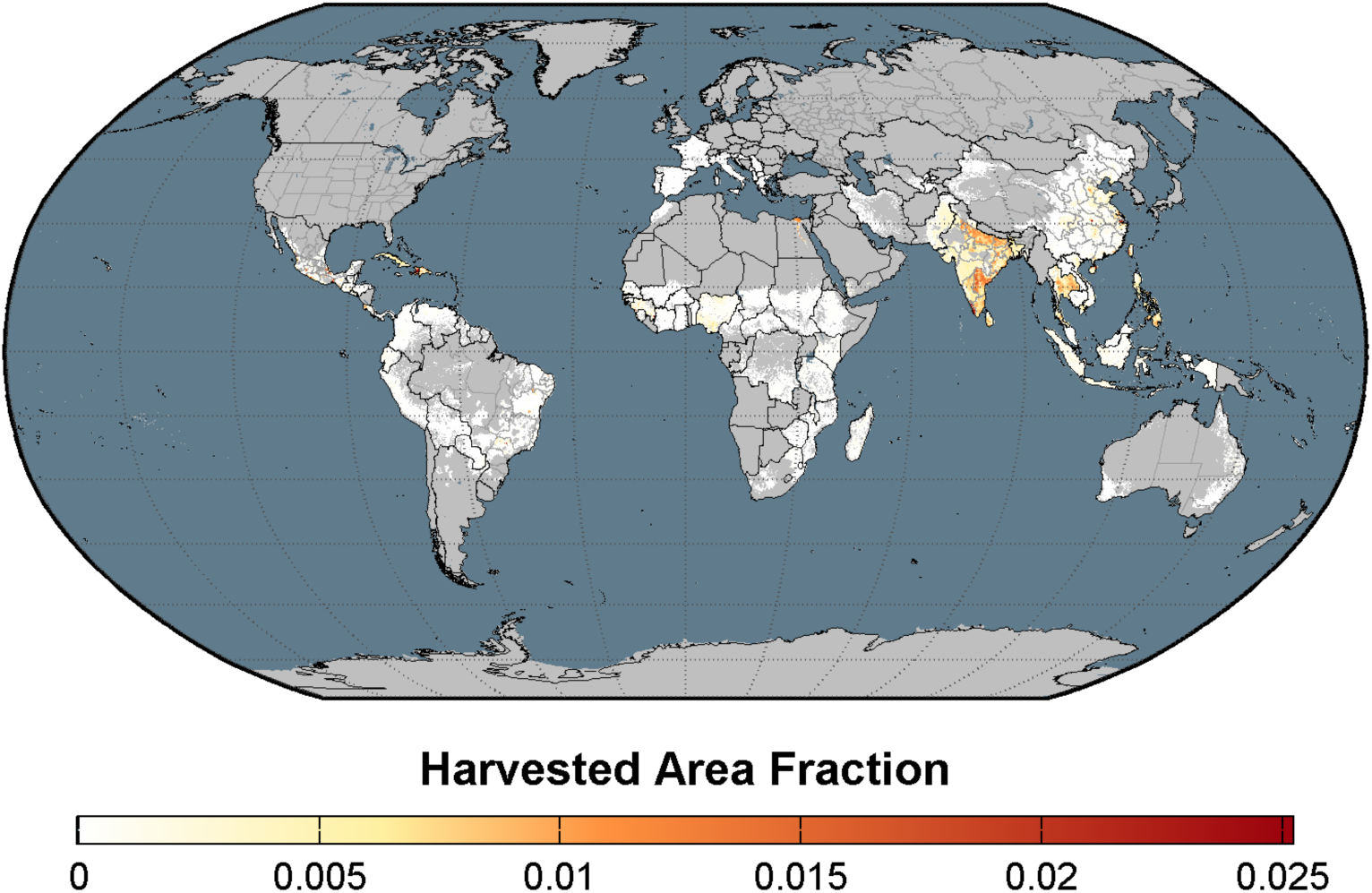
Harvested Area Fraction of domesticated mango (Mangifera indica). Data adapted from Monfreda et al. (2008) by EarthStat.org.

**Supplementary Figure II:**
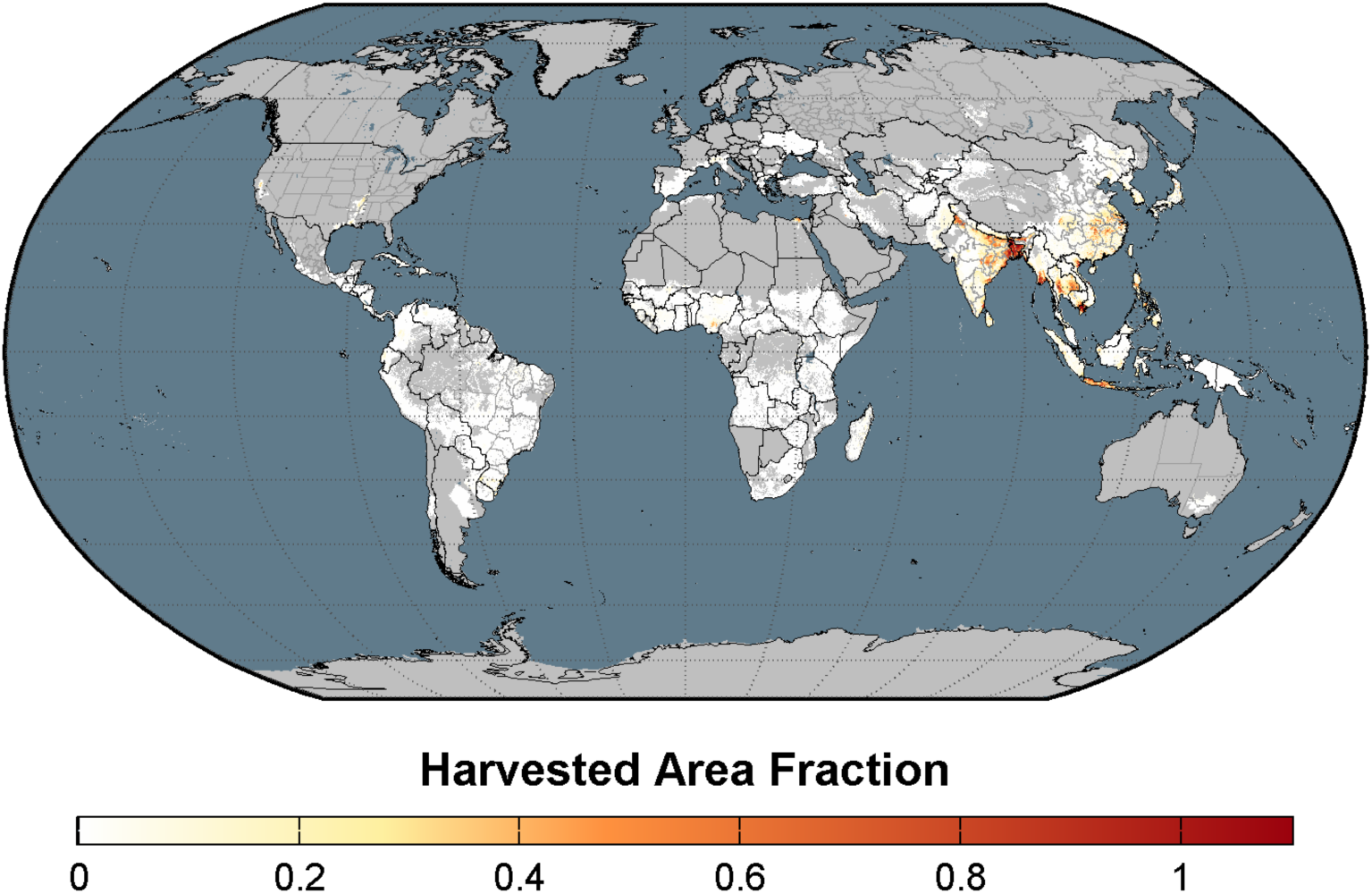
Harvested Area Fraction of domesticated rice (Oryza sativa). Data adapted from Monfreda et al. (2008) by EarthStat.org.

**Supplementary Figure III:**
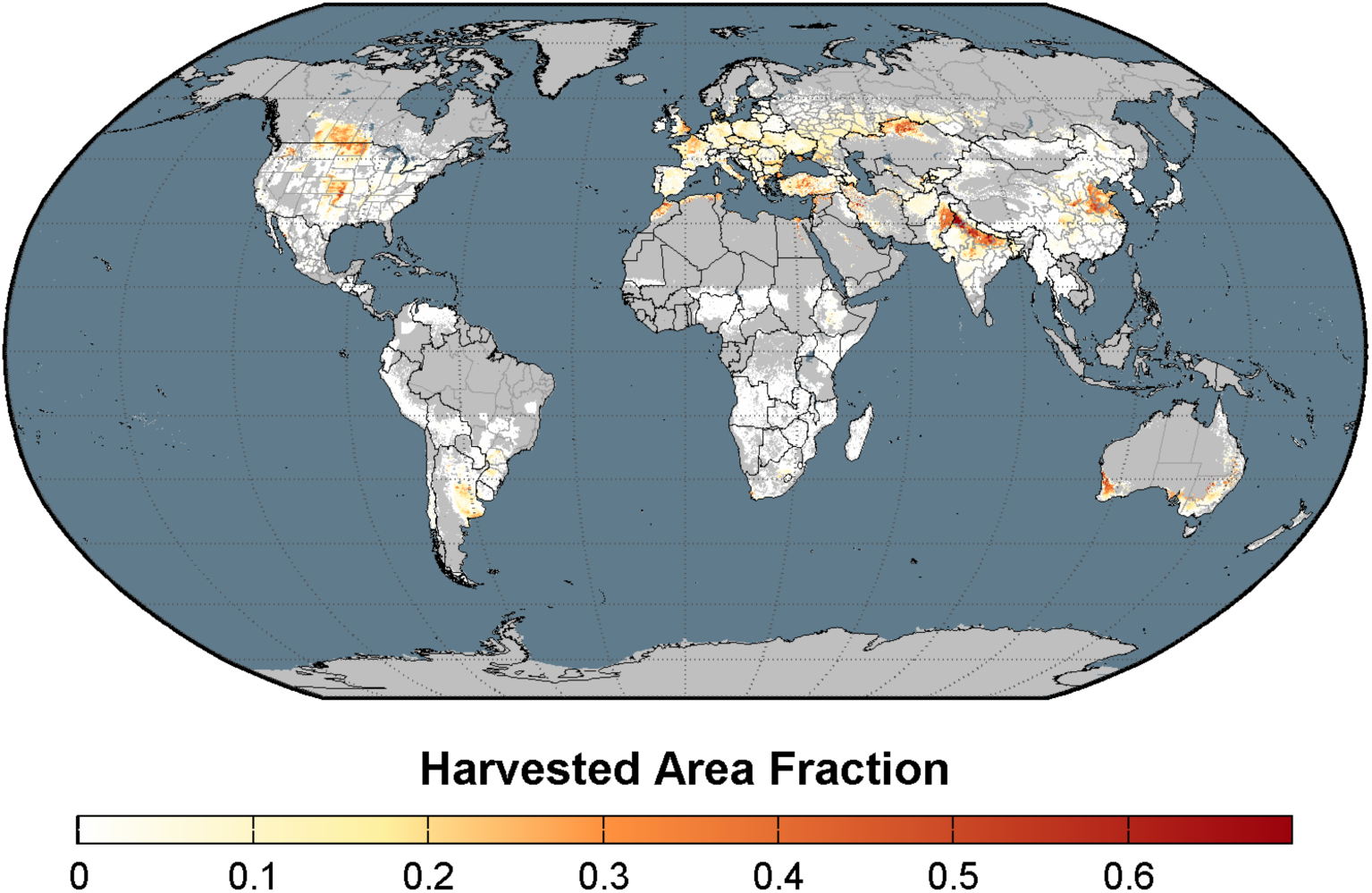
Harvested Area Fraction of domesticated (common) wheat (Triticum aestivum). Data adapted from Monfreda et al. (2008) by EarthStat.org.

**Supplementary Figure IV:**
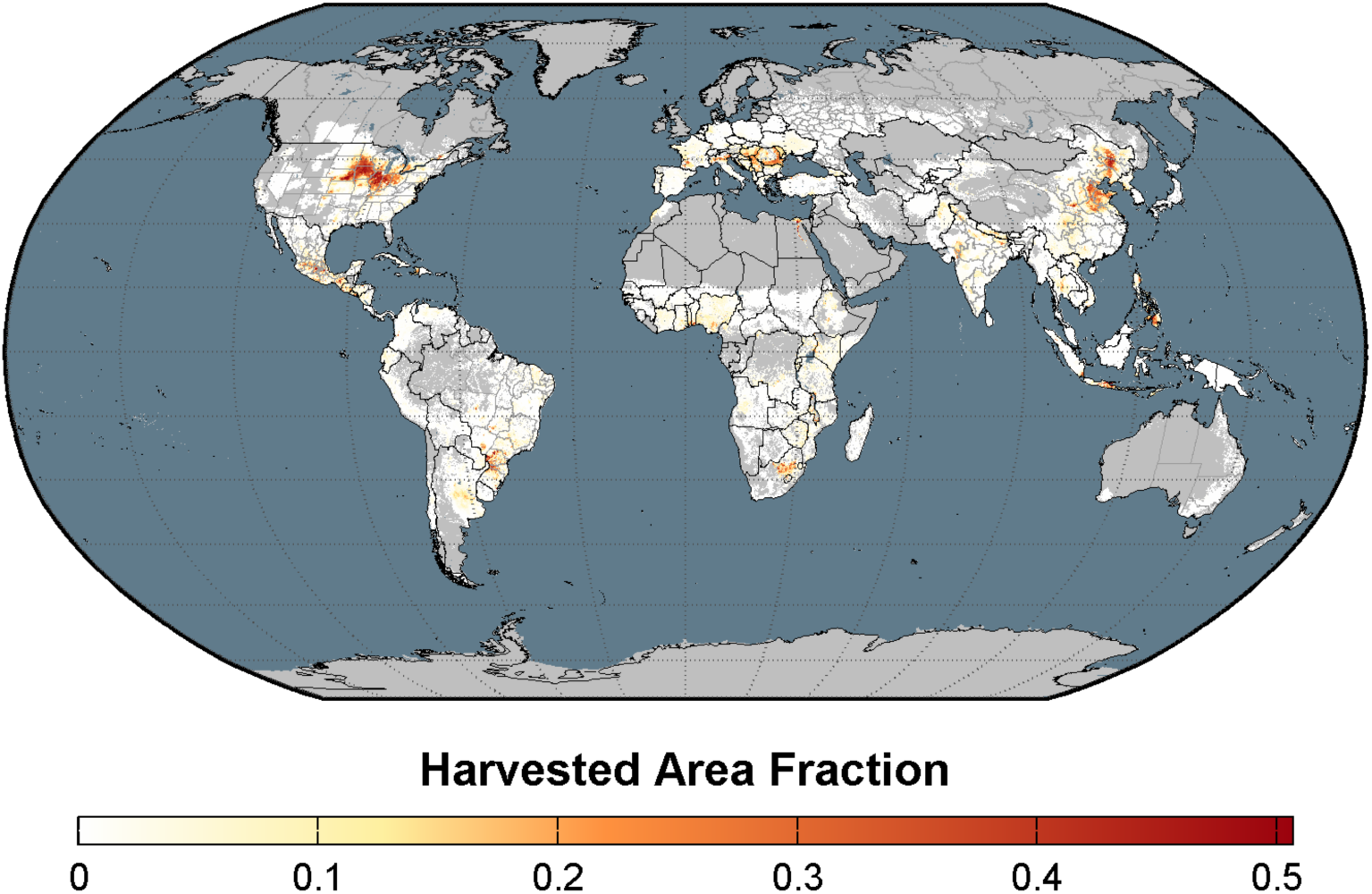
Harvested Area Fraction for domesticated maize (Zea mays). Data adapted from Monfreda et al. (2008) by EarthStat.org.

